# Library-based analysis reveals segment and length dependent characteristics of defective influenza genomes

**DOI:** 10.1101/2021.09.06.459165

**Authors:** Marisa Mendes, Alistair B. Russell

## Abstract

Parasitic elements of the viral population which are unable to replicate on their own yet rise to high frequencies, defective interfering particles are found in a variety of different viruses. Their presence is associated with a loss of population fitness, both through the depletion of key cellular resources and the stimulation of innate immunity. For influenza A virus, these particles contain large internal deletions in the genomic segments which encode components of the heterotrimeric polymerase. Using a library-based approach, we comprehensively profile the growth and replication of defective influenza species, demonstrating that they possess an advantage during genome replication, and that exclusion during packaging reshapes population composition in a manner consistent with their final, observed, distribution in natural populations. We find that an innate immune response is not linked to the size of a deletion; however, replication of defective segments can enhance their immunostimulatory properties. Overall, our results address several key questions in defective influenza A virus biology, and the methods we have developed to answer those questions may be broadly applied to other defective viruses.

## Introduction

It has long been known that the total number of particles in a viral population far exceeds the number of infectious particles (***Donald and Isaacs, 1954***).

Early work on this phenomenon revealed that, in addition to fully-infectious particles, there are semi-infectious, interferon-suppressive, and defective-interfering particles (DIPs) (***Brooke, 2014***; ***Ngunjiri et al., 2008***; ***Marcus et al., 2005***; ***Brooke et al., 2013***; ***Akpinar et al., 2016***). The latter is marked by their capacity to inhibit, rather than enhance, infection. Interference by DIPs can arise from direct competition for resources within an infected cell, exclusion of replication-competent viruses during packaging, or increasing the production of interferons, key signaling components of cell-intrinsic innate immunity (***Genoyer and López, 2019***; ***Tapia et al., 2013***; ***Odagiri and Tashiro, 1997***; ***Laske et al., 2016***). While the genetic basis of DIPs is idiosyncratic, differing from virus to virus, a defining characteristic is their capacity to, despite their defects, rise to high frequencies in viral populations when complemented by wild-type virus in coinfections.

Due to their association with coinfections, DIPs were once largely thought to be artifacts of laboratory-grown viruses (***Vignuzzi and López, 2019***). However, clinical sequencing has revealed that they are a common component of viral infections and can be associated with disease outcome (***Felt et al., 2021***; ***Saira et al., 2013***; ***Vasilijevic et al., 2017***). Their potential ability to modulate the course of disease has now sparked a renewed interest in their therapeutic potential (***Rezelj et al., 2021***; ***Rand et al., 2021***; ***Tanner et al., 2019***; ***Dimmock and Easton, 2014***).

For influenza A virus, a (-) sense, segmented, RNA virus, defective particles, interfering or otherwise, largely bear genomes with large internal deletions in the three polymerase-encoding segments (***Nayak et al., 1982***; ***Nakajima et al., 1979***; ***Akkina et al., 1984***; ***Alnaji et al., 2019***). These deletions generally reduce the size of each segment from ∼2.3kb to ∼400nt (***Saira et al., 2013***; ***Pelz et al., 2021***). It is unknown what drives this size and segment preference. Understanding deletions, from formation through selection, is challenging. Studying *de novo* deletion formation, a stochastic event, requires single-cell resolution, as was performed by Kupke *et al*, or a highly sensitive sequencing approach in order to disambiguate rare production of deletions from artifacts such as template-switching (***Kupke et al., 2020***; ***Cocquet et al., 2006***). Even with those controls, observed deletions remain subject to selective pressures from genome replication. Rather than focus on their formation, and the challenges that presents, we instead chose to study selection on deletions *after* they form in order to understand if these pressures may nevetheless provide an understanding of the ecology of these particles.

Using a combination of PCR and Gibson assembly we seeded an initial population with a variety of deletions and duplications that generated a broad range of lengths, and measured changes in this distribution as a function of progression through the viral life cycle. While validated full-length, amplicon-based, approaches exist for sequencing influenza A virus, they exhibit length-dependent biases, which can artifactually generate apparent differences in deletion abundance due to differences in saturation kinetics (***Boussier et al., 2020***). To overcome this limitation, we instead used an approach wherein individual barcoded adapters were associated with a particular deletion, or duplication. Our libraries were similar to those generated using RanDeL-Seq, a recent effort wherein deletions were generated in HIV and Zika genomes using random transposon insertions and exonuclease digestion (***Notton et al., 2021***).

Using our artificial libraries, we demonstrate that deletions in influenza segments lead to significant advantages during genome replication, explaining their abundance. Certain deletions are excluded during packaging, both in terms of size and segment identity, likely explaining the observed distribution in final viral populations. Lastly, we find that deletions can generate aberrant sequences that can drive interferon induction, but their stimulatory behavior is not length-dependent, and is enhanced by genome replication.

## Results

### PCR-based generation of length polymorphic deletion/duplication libraries

The observed frequencies of variants within an initially clonal population are not only reflective of the fitness benefits, or drawbacks, of any given variant, but also the timing with which they are formed (***Luria and Delbrück, 1943***). In other words, the distribution of defective segments is highly confounded by whether a deletion forms early in population expansion or late. To better understand how defective influenza species may compete amongst one-another, we designed a library-based approach wherein we introduce, contemporaneously, a distribution of more than 100 possible lengths representing over 1,000 possible junction combinations into an influenza genomic segment. We thereafter track the distribution of variants within each library as a function of progression through the viral lifecycle. Our library approach is described in (***Figure 1***A).

**Figure 1.**
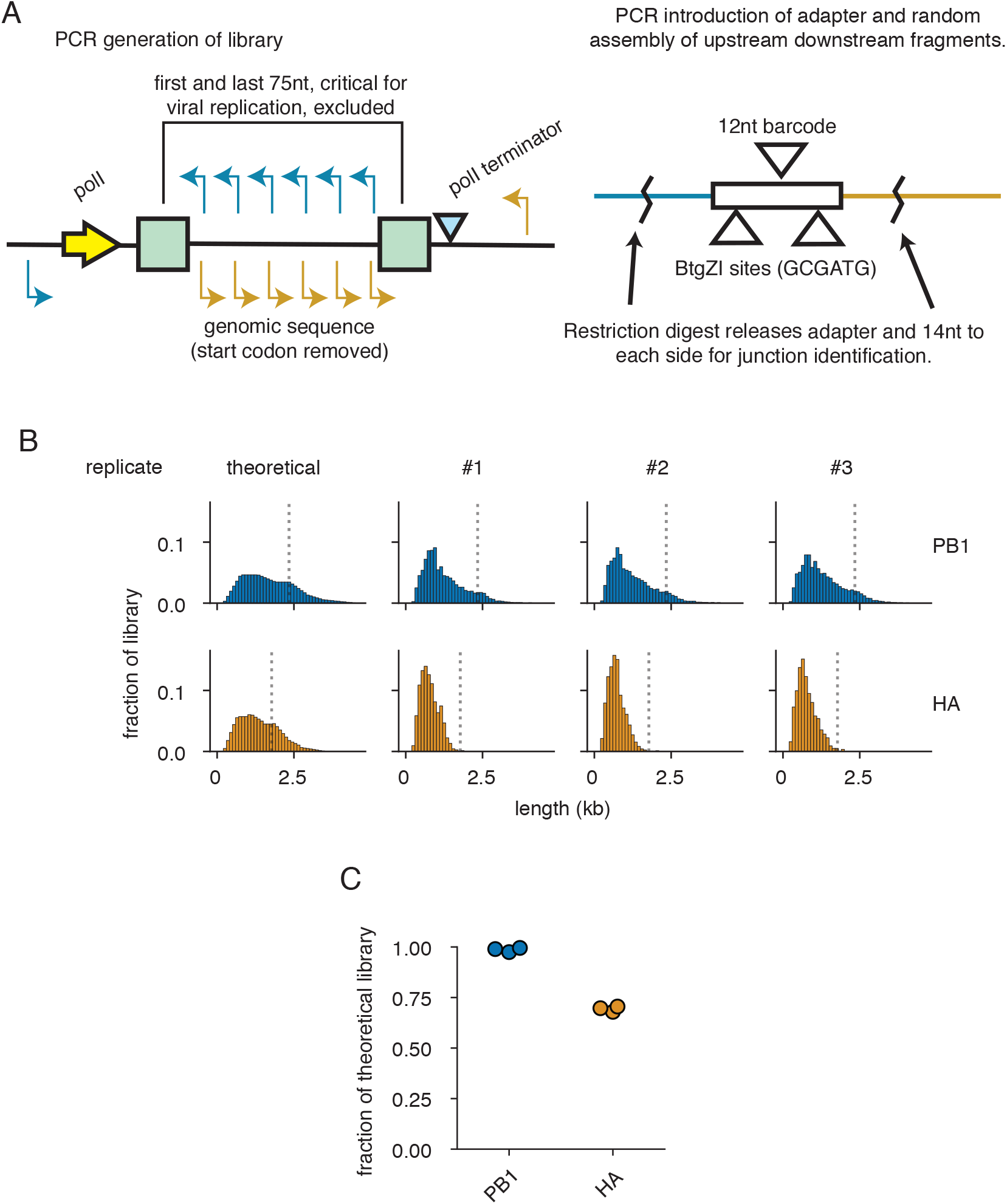
A PCR-based strategy for exploring length-dependent phenomena in viral libraries. **(A)** As templates for each library, we used vectors encoding each segment under control of a polymerase I-dependent system for generating authentic viral RNAs (***Neumann et al., 1999***). For each library, a set of staggered primers were used in individual PCR reactions with an external primer. Further PCR was used to append a 57nt adapter with a random, 12nt, barcode. Products were then randomly combined in a pooled Gibson assembly reaction. To link barcodes to duplication or deletion junctions, barcodes and flanking regions were freed from the plasmid pool with a BtgZI digest and sequenced. **(B)** Theoretical and actual length distributions of libraries. Lengths do not include the 57nt adapter. Bins in this and all other panels are 100nt in length, numbers represent the right edge of each bin. Dotted lines show the length of wild-type PB1 and HA. **(C)** Libraries contain most of the theoretical diversity. Each point represents a single library. **Figure 1–source data 1**. primers used to generate the library are in `libraryPrimers.tsv`.

In brief, we computationally designed primers in both the forward and reverse orientations across target influenza segments staggered every 25nt for the first third of the segment, 50nt for the second, and 100nt for the last. The first and last 75nt of each segment were excluded as they contain the viral promoter and critical packaging sequences (***Odagiri and Tashiro, 1997***; ***Liang et al., 2005***; ***Goto et al., 2013***; ***Azzeh et al., 2001***; ***Flick et al., 1996***). Each individual primer with the same orientation was paired with a specific, shared, primer, and individual PCRs were performed to generate forward and reverse flanks of staggered sizes. An additional round of PCR was performed on reverse flanks to add a 57bp adapter containing a 12bp random barcode, flanked by cut sites for the type IIS restriction enzyme BtgZI. Gibson assembly was then used to randomly pair forward and reverse flanks, reconstructing a library consisting of both deletions and duplications (***Gibson et al., 2009***). Taking advantage of the capacity of type IIS restriction enzymes to cleave outside of their restriction sites, we released barcoded adapters with 14/10nt of flanking sequence with a BtgZI digest. By sequencing these adapters, we may link specific deletions, or duplications, to their corresponding barcode. Thereafter, to infer junction abundance we can sequence across our variable barcodes from constant regions in our adapter. Our barcoding schema avoids the length-bias of target amplicon approaches to sequencing influenza, which overestimate the abundance of shorter species relative to longer (***Boussier et al., 2020***).

Influenza A virus defective particles largely consist of those bearing large deletions in the three polymerase-encoding segments, as deletions in the remaining five segments appear only rarely in viral populations (***Nayak et al., 1982***; ***Nakajima et al., 1979***; ***Akkina et al., 1984***; ***Alnaji et al., 2019***). There is currently no single explanation for this observation. Possibilities include differences between segments in the rate of deletion formation, replication, or even packaging (***DUHAUT and MCCAULEY, 1996***; ***Laske et al., 2016***). To explore whether features other than biases in formation may contribute to the observed differences in abundance, we chose to generate libraries in a single polymerase segment, PB1, encoding the core viral polymerase, and a single non-polymerase segment, HA, encoding the viral receptor-binding and fusion protein hemagglutinin. In order to focus solely on length and sequence composition, and not on whether a given segment produces a functional protein, we used variants of PB1 and HA with mutated start codons. Using the schema described above, we expect 1,936 possible unique junctions for PB1 and 1,225 possible junctions for HA. After library generation, we find a slight bias towards shorter lengths, potentially due to higher molar yield in PCR reactions, and relatively complete representation of unique junctions (∼98% of theoretical PB1 junctions and ∼70% of theoretical HA junctions) (***Figure 1***B, C).

### Shorter segments are preferentially amplified during genome replication

The first question we wished to address with these libraries was whether length, independent of other features of the viral life cycle, influences genome replication. There are conflicting reports concerning whether shorter segments replicate faster, and thus outcompete their longer counterparts (***Laske et al., 2016***; ***Alnaji et al., 2021***). To make these measurements, we co-transfected libraries with plasmids encoding the minimal influenza genome replication machinery—PB2, PB1, PA, and NP—and measured barcode abundance in viral genomes after 24 hours. To determine enrichment, or depletion, of a given barcode, we generated an orthogonal dataset consisting of the same plasmid libraries transfected with PB1, PA, and NP, but not PB2. By generating a replication-incompetent dataset, we corrected for length-dependent transcription and degradation biases from the polymerase I-dependent viral RNA transcription system (***Figure 2–Figure Supplement 1***) (***Neumann et al., 1999***).

Comparing the composition of libraries between replication-incompetent, and -competent, conditions, there is a noticeable shift towards smaller variants (***Figure 2***A). Plotting enrichment as a function of length, these two features are negatively correlated, with Spearman correlation coefficients of -0.86 and -0.92 for PB1 and HA respectively (***Figure 2***B). Smaller segments that arise due to deletions would therefore be expected to possess a considerable replication advantage over their longer counterparts. This observation is in line with prior work from Widjaja *et al*. showing shorter, Gaussia, luciferase was preferentially replicated by viral replication machinery over longer, firefly, luciferase (***Widjaja et al., 2012***). To confirm this conclusion, we generated PB1 and HA segments of defined length (200, 400, 800, and 1600nt) consisting of equal numbers of nucleotides from the 5’ and 3’ termini of each segment. As in our libraries, we introduced an additional 57nt adapter with a barcode; however, rather than a random barcode we instead used a defined barcode to delineate each size. A control qPCR demonstrates our ability to disambiguate each of these barcodes from one-another, with greater than 1000-fold discrimination for target versus off-target amplification (***Figure 1–Figure Supplement 2***). As expected, there remains a clear inverse relationship between enrichment in a genome replication assay and length for both segments at a Spearman correlation coefficient of -0.95 for both PB1 and HA (***Figure 2***C). Therefore, at least in part, a length-dependent replication advantage explains how these parasitic, deleterious, variants rise to frequencies exceeding their longer, replication-competent, progenitors. Nevertheless, we procure no explanation for why deletions are more frequently observed in PB1 than HA.

**Figure 2.**
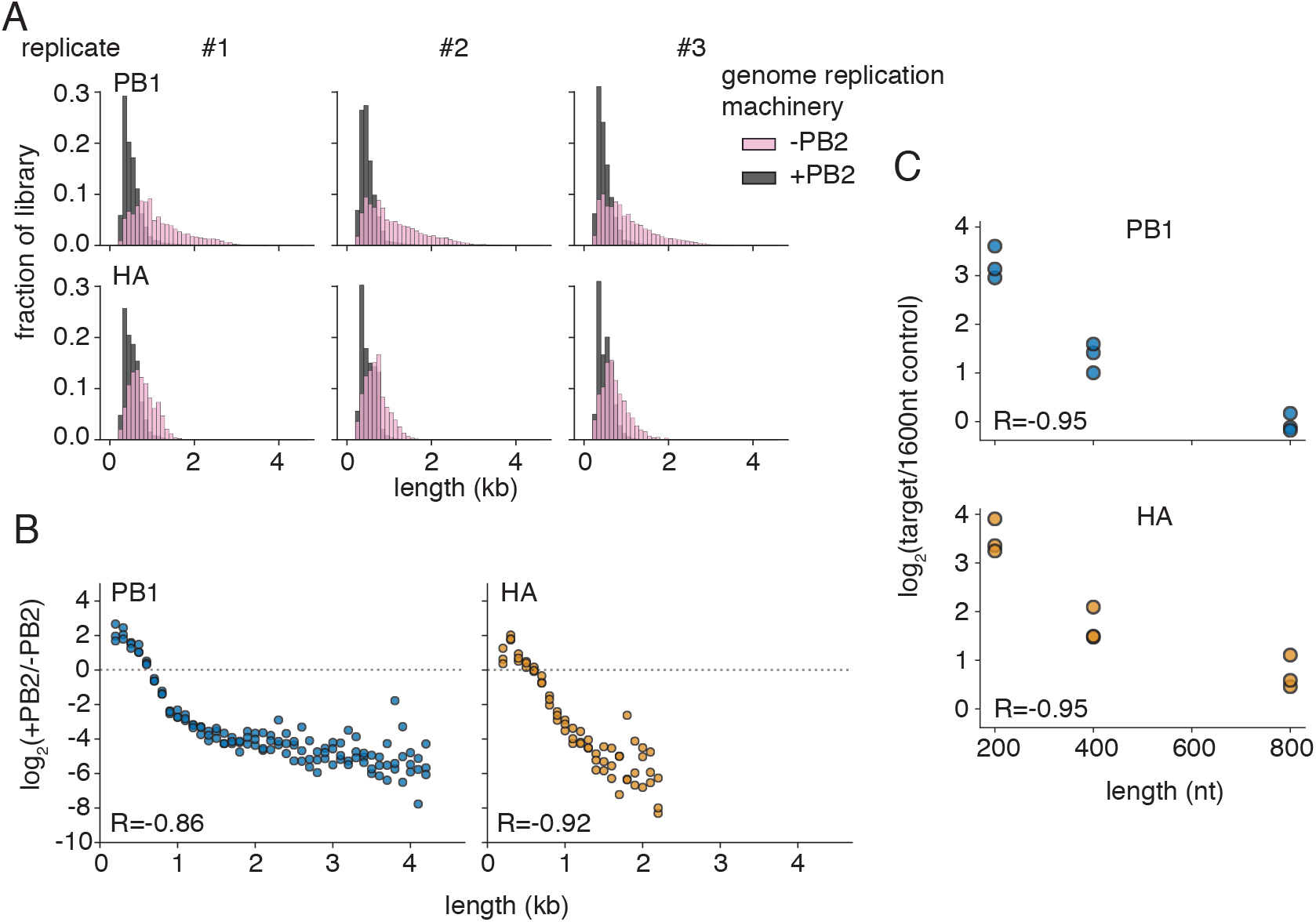
Segment length and genome replication efficiency are inversely correlated. **(A)** Size distributions of libraries in transfections with (+PB2) and without (-PB2) the full viral genome replication machinery 24 hours post-transfection in 293t cells. **(B)** Smaller species are enriched during genome replication. The fraction of variants falling within each 100nt bin was compared between replication-competent (+PB2), and -incompetent (-PB2) selections. Points above the dotted line represent sizes in each individual library which were enriched in the +PB2 dataset. Points were only shown if represented in all three libraries under both conditions R is the Spearman correlation coefficient. n=3. **(C)** qPCR supports the conclusion that genome replication is inversely correlated with length. Vectors encoding variants of four seperate lengths (200, 400, 800, and 1600nt) were transfected into 293t cells with and without the full viral genome replication machinery and analyzed by qPCR 24 hours post transfection. The fraction of each variant was calculated relative to the longest variant (1600nt), and this value compared between replication-competent and -incompetent datasets to determine the relative enrichment attributable to genome replication. R is the Spearman correlation coefficient. n=3. **Figure 2–Figure supplement 1**. Polymerase I length biases, related to ***Figure 2***A **Figure 2–Figure supplement 2**. Validation of qPCR in ***Figure 2***C.

### PB1 and HA exhibit length-dependent differences in viral incorporation

In addition to an advantage during genome replication, it has also been posited that segments bearing deletions might outcompete their full-length counterparts in packaging, although recent work has suggested that the opposite may be true (***DUHAUT and MCCAULEY, 1996***; ***Alnaji et al., 2021***). To explore aspects of the viral lifecycle uncaptured in our genome replication assay, we transfected cells with our libraries and the viral replication machinery and waited 24 hours. Thereafter, we infected these transfected cells with wild-type virus and allowed infection to proceed for 72 hours. During this time, variants within each population could be complemented by coinfection with wild-type virus and compete with one-another for packaging into viral particles. We then harvested clarified supernatant and sequenced those variants that had become incorporated into the viral population.

Comparing the viral supernatant to replication-only controls, there are divergent patterns of enrichment between HA and PB1 libraries (***Figure 3***A). For HA, there is a direct, positive, relationship between length and enrichment in the viral supernatant, with considerable advantage to longer, over shorter, segments (Spearman correlation coefficient of 0.94) (***Figure 3***B). This is consistent with recent work showing disperse packaging signals—the more HA sequence a variant contains the better the capacity for the segment to become incorporated into viral particles (***Dadonaite et al., 2019***; ***Sage et al., 2020***; ***Marsh et al., 2007***). For PB1, small segments, <400nt of natural sequence, are excluded but otherwise preferences are relatively flat. This suggests that additional PB1 sequence beyond the first 400nt provides little benefit, indicating that packaging requirements for PB1 are divergent from those for HA. This is reflected in a much more modest correlation between length and enrichment at a Spearman correlation coefficient of 0.55. To confirm that PB1 and HA exhibit divergent relationships between length and packaging, we used defined 400nt and 1600nt variants and repeated selection as performed for our more extensive libraries. As expected, qPCR analysis finds more robust enrichment of a 1600nt length segment over a 400nt counterpart in HA than in PB1 (***Figure 3***C). Therefore divergent kinetics in incorporation into the viral population, likely due to packaging requirements, provide an explanation for why deletions are more rarely observed in HA than for PB1.

**Figure 3.**
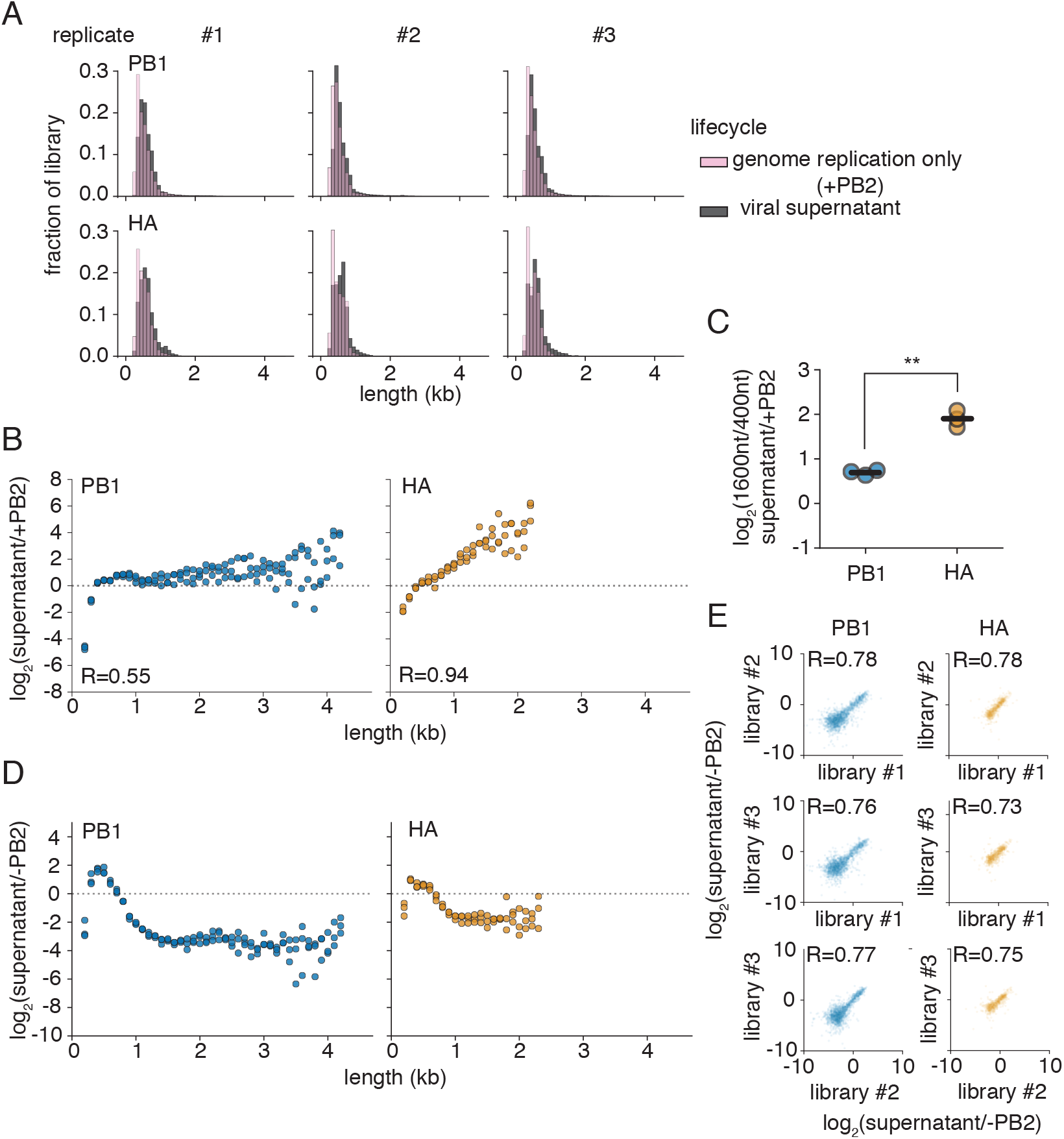
PB1 exhibits a reduced length-dependence on packaging relative to HA. **(A)** Variant distribution after replication alone (+PB2), or in the supernatant after 72h of infection with wild-type, virus. **(B)** Packaging is strongly correlated with length for the HA segment. The fraction of variants falling within each 100nt bin was compared between replication only, and supernatant selections. Lengths above the dotted line are over-represented in packaged viral particles relative to a replication only control. Points were only shown if represented in all three libraries under both conditions R is the Spearman correlation coefficient. n=3. **(C)** qPCR supports the conclusion that PB1 and HA exhibit divergent behaviors in packaging. The ratio of a 1600nt to a 400nt product was compared between replication only and supernatant samples as in **(B)**. Higher numbers indicate preferential packaging of a 1600nt product in the supernatant. Each point is a single replicate, lines are the mean. Asterisks indicate p<0.05, two-tailed t-test. n=3 **(D)** Conflicting pressures of genome replication shape defective populations. The fraction of variants falling within each 100nt bin was compared between replication-incompetent (-PB2), and viral supernatant selections. Lengths above the dotted line are overrepresented in the supernatant compared to their original presence in the polI-transcribed library. Points were only shown if represented in all three libraries under both conditions. n=3. **(E)** Data in **(D)** are highly replicable between libraries. Inter-replicate enrichment values as calculated in **(D)**. R is the Pearson correlation coefficient. **Figure 3–Figure supplement 1**. Length distributions for ***Figure 3***D.

Assaying the entire process, from polI transcription to successful encapsidation in viral particles in the supernatant, we see the conflicting pressures of replication efficiency and incorporation into the viral population produce a highly stereotyped deletion enrichment curve for both PB1 and HA, with maxima around 400nt in length (***Figure 3***D). Deletions with a final length centering around this size become enriched in our library, whereas those that are smaller or larger become depleted. This process was highly replicable between independently-generated libraries with inter-replicate Pearson correlation coefficients of ∼0.75 when comparing across the enrichment or depletion of individual junctions (***Figure 3***E). These maxima correlate well with previously-identified “sweet-spot” lengths from continuous culture propagation of influenza defective particles (***Pelz et al., 2021***). Taken together, our data indicate that the length distribution of defective influenza segments can be explained by the competing processes of replication and packaging, regardless of their initial distribution upon formation.

### Deletions in the influenza genome are associated with interferon-producing cells

The work of Tapia *et al*., since repeated by others, demonstrated that populations replete in defective influenza particles exhibit enhanced interferon induction (***Tapia et al., 2013***; ***Liu et al., 2019***). To explore further, we wished to probe whether our libraries capture the variation responsible for this increase in the innate immune response. Unlike our previous selections, where viral replication and propagation served to shape variant distribution, understanding how variants may influence innate immune detection is less straightfoward. Potentially solving this problem, prior work indicates that observed increases in interferon induction in response to influenza variants are frequently due to more cells producing interferon rather than an increase in the amount of interferon each infected cell produces (***Chen et al., 2010***; ***Killip et al., 2017***; ***Russell et al., 2019***). As such, if we sort infected cells based on interferon production, we would anticipate that we may be able to co-enrich stimulatory variants in our libraries.

Before applying this selection to our libraries, we first sought to understand whether it could recapitulate the earlier findings that deletions broadly associate with interferon induction. To this end, we first generated two wild-type influenza A virus populations that contained an intermediate burden of defective particles, independently, from reverse-genetics onwards (***Figure 4–Figure Supplement 1***). By working with a population where defective particles are neither rare nor overwhelming, we allow the potential for enrichment of deletions in interferon-producing cells; otherwise, if all cells are infected by defective viruses we may be unable to achieve any additional enrichment, and if few to none are we may lack the sensitivity to detect their presence. We next infected an A549 cell line expressing the cell-surface marker LNGFR under control of the *IFNB1* promoter and sorted populations into interferon-enriched, and -depleted, subsets using magnetic-activated cell sorting (***Russell et al., 2019***). Prior to sequencing viral genomes from these populations, we first validated our sorting efficiency with total mRNA sequencing of both populations (***Figure 4***A). As expected, there is significant enrichment of the gene encoding the most highly-transcribed type I interferon in A549 cells, interferon beta (*IFNB1*). We also find co-enrichment of type III interferon transcripts (*IFNL1, IFNL2*, and *IFNL3*). Additional enriched genes include a number of interferon-stimulated genes, suggesting either direct transcription of a subset of ISG loci by IRF3, or a high level of autocrine signaling experienced by interferon-producing cells.

**Figure 4.**
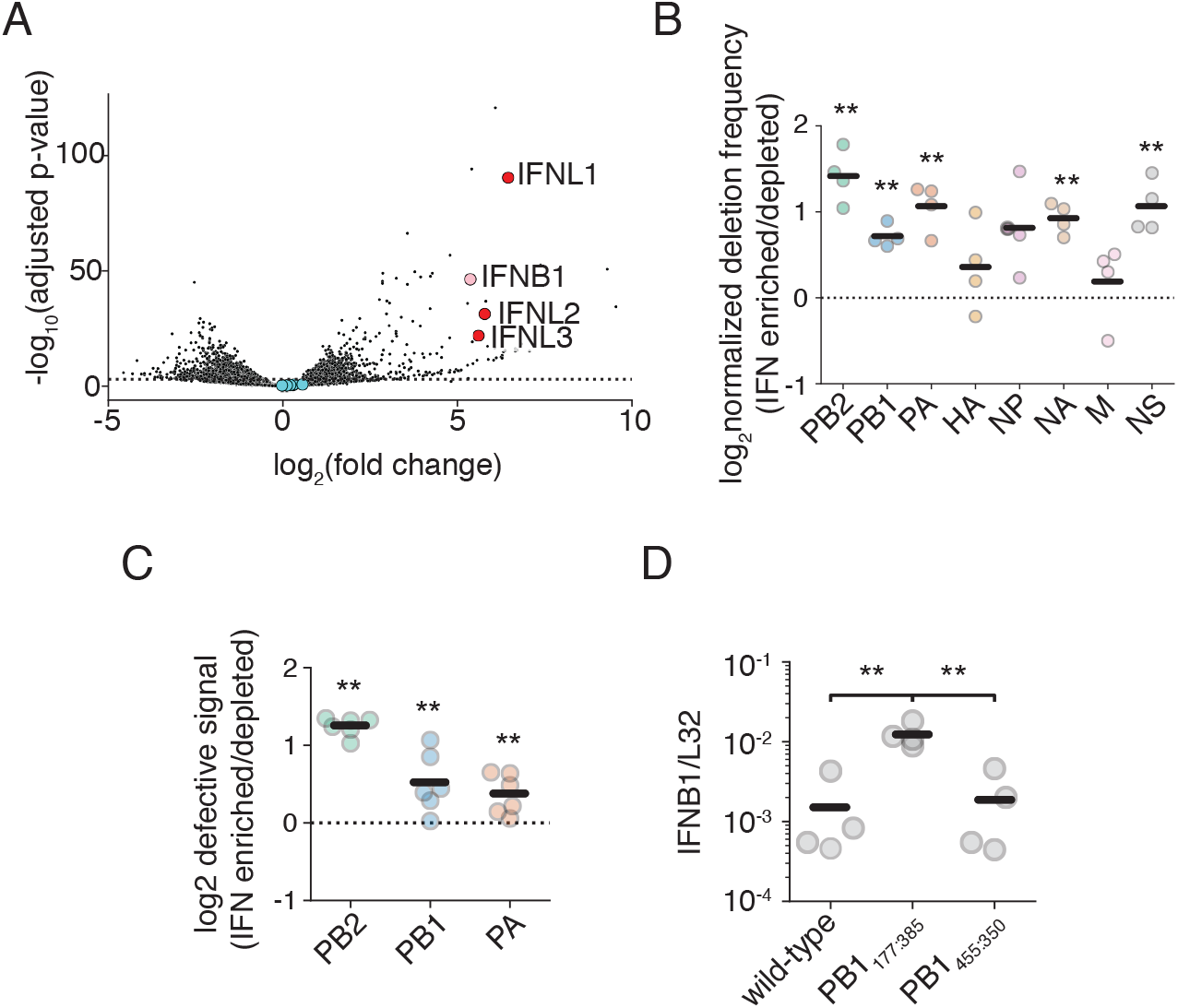
Interferon-producing cells are enriched for defective influenza segments. **(A)** Differential gene expression analysis on interferon-enriched and -depleted populations. An A549 type I interferon reporter cell line was infected at an MOI of 0.1 and cells were sorted using magnetic-activated cell sorting 14h post infection. Fold-change calculated as mRNA abundance in interferon-enriched/interferon-depleted datasets. Type I interferon gene *IFNB1* labeled in pink, the type III interferon genes *IFNL1-3* in red, and influenza transcripts in blue. n=4. **(B)** The fraction of junction-spanning fragments per influenza segment was determined for both interferon-enriched and depleted populations, and compared. Values above the dotted line indicate segments with deletions that co-enriched with inteferon. Asterisks represent significant enrichment, one-sample one-tailed t-test. n=4. **(C)** qPCR validation of **(B)** for the three polymerase segments. A qPCR responsive to defective burden as described in ***Figure 4–Figure Supplement 6*** was used. Values above the dotted line indicate a co-enrichment of deletions with interferon. Asterisks represent significant enrichment, one-sample one-tailed t-test. Triplicate measurements were made of two biological replicates. n=6. **(D)** qPCR measuring levels of *IFNB1* transcripts normalized to the house-keeping control *L32* 8h post infection from A549 cells infected at an MOI of 0.5. Asterisks represent significantly different values in all pairwise comparisons, two-tailed t-test. n=4. For panels **(B), (C)**, and **(D)** individual points are individual measurements, lines are mean values. For panels **(B), (C)**, and **(D)** siginficance was calculated using Benjamini-Hochberg correction at an FDR of 0.05. **Figure 4–Figure supplement 1**. Confirmation of defective burden of populations used in ***Figure 4***A, B, and C. **Figure 4–Figure supplement 2**. Sampling stochasticity for interferon-enriched samples for (***Figure 4***B). **Figure 4–Figure supplement 3**. Sampling stochasticity for interferon-depleted samples for ***Figure 4***B. **Figure 4–Figure supplement 4**. Raw data for ***Figure 4***B. **Figure 4–Figure supplement 5**. Mapping depth for data from ***Figure 4***B. **Figure 4–Figure supplement 6**. Validation of qPCR in ***Figure 4***C. **Figure 4–Figure supplement 7**. Validation of infections in ***Figure 4***D. **Figure 4–source data 1**. full differential gene expression data are in `DESeq2.tsv`.

We next used a full-length PCR-based strategy to sequence influenza genomes (***Hoffmann et al., 2001***). While length-biased, we sought to reduce this bias by initiating reverse transcription with equivalent viral RNA, and limiting cycles of PCR. We identified reads containing deletion junctions by mapping continuous reads with STAR, identifying discontinuous mapping in unmapped reads with BLAST, collapsing all identified junctions into a single annotation file for each biological replicate, and then remapping discontinuous reads using this BLAST-annotated STAR index (***Dobin et al., 2013***; ***Altschul et al., 1990***). We required that a deletion had three mapped bases to either side of a junction and that read pairmates were concordant. As we used paired-end reading, deletion junction counts were collapsed to a per fragment value rather than per read value. As expected from the highly stochastic nature of deletion formation, while junction counts between infection replicates using the same biological population are highly correlated, those from divergent biological replicates exhibit no detectable correlation to one-another (***Figure 4–Figure Supplement 2, Figure 4–Figure Supplement 3***).

Deletion-spanning fragments are predominantly derived from the polymerase segments, regardless of whether a sample was enriched or depleted for interferon (***Figure 4–Figure Supplement 4***). When we compare the fraction of deletion-containing fragments between interferon-enriched and depleted datasets, there is statistically significant enrichment across most segments (***Figure 4***B). Mapping depth across the polymerase segments is consistent with this enrichment, showing a slight, but replicable, increase in depth at the 5’ and 3’ ends of the three polymerase segments relative to the central portion of each gene segment (***Figure 4–Figure Supplement 5***). Other deletions, while enriched, were too rare to produce visible changes in sequencing depth. To confirm these results were not driven by PCR bias, we developed a qPCR sensitive to increased defective content (***Figure 4–Figure Supplement 6***). Our design leverages the short extension time common in most qPCR protocols, and so, by designing primers flanking the shared limits of deletion breakpoints between biological replicates we generate a signal that is responsive to the presence of shortened products while producing little to no signal on full-length polymerase segments. We can then correct this gestalt “DI-specific” signal to “full-length” signal generated from qPCR primers that lie nested within most observed deletions (***Schwartz and Lowen, 2016***). Using this qPCR, we confirm our deep-sequencing results (***Figure 4***C).

As deletions preferentially associate with interferon induction, we may ask whether all deletions exhibit an identical relationship to this process. That is, is the mere presence of a deletion sufficient to induce high levels of interferon, or do different deletions exhibit differential induction of interferon pathways? Using complementing cells, we grew a previously-cloned stimulatory PB1 deletion containing 177nt from the 5’ end of the vRNA, and 385nt from the 3’ end of the vRNA, PB1_177:385_, and a newly-identified deletion from our dataset, PB1_455:350_ (***Russell et al., 2019***). Comparing the two, we find that the former retains its capacity in infections to induce high leves of type I interferon transcripts, whereas the latter is no more stimulatory than a nominally wild-type population (***Figure 4***D).

### Interferon induction by defective segments is a length-independent process

As different deletions can cause divergent levels of interferon stimulation, and as we may sort by interferon production to classify immunostimulatory variation, we may use our sortable cell lines and our libraries to generate comprehensive measurements of how length of a defective segment might impact interferon induction. The increase in interferon observed in infections with defective influenza species has been posited to arise from a preference for shorter molecular species by RIG-I, the major sensor of this virus (***Rehwinkel et al., 2010***; ***Baum et al., 2010***). However, recent experiments found that while shorter influenza species do exhibit enhanced interferon stimulation, the size at which this occurs, <125nt, is far smaller than defective influenza genomes (***Velthuis et al., 2018b***). To determine whether, like with packaging and genome replication, interferon induction is also a length-dependent process in our libraries, we infected a type III interferon reporter A549 cell line with our libraries and sorted on interferon production.

We changed to a cell line expressing LNGFR and eGFP under control of a type III interferon promoter over a type I reporter as the sort described in ***Figure 4*** only produced a yield of ∼5000 cells per sort from a starting population of 20 million cells. As the theoretical yield based on the fraction of interferon-positive cells ranges from ∼100,000-200,000, this represents a greater-than 80% loss of material, and would produce far too low a yield to analyze a comprehensive set of variants. Based on our mRNAseq data, and the prior observation that a type III interferon reporter line produces a stronger signal that still correlates with type I interferon production, we opted to change strategies (***Russell et al., 2019***). Consistent with an improved resolution, we were able to recover ∼100,000 cells in each of our interferon-enriched sorts, and, by qPCR, our sorting efficiency as measured by type I interferon expression was comparable to that achieved with a type I interferon reporter (***Figure 5***A, ***Figure 5–Figure Supplement 1***).

**Figure 5.**
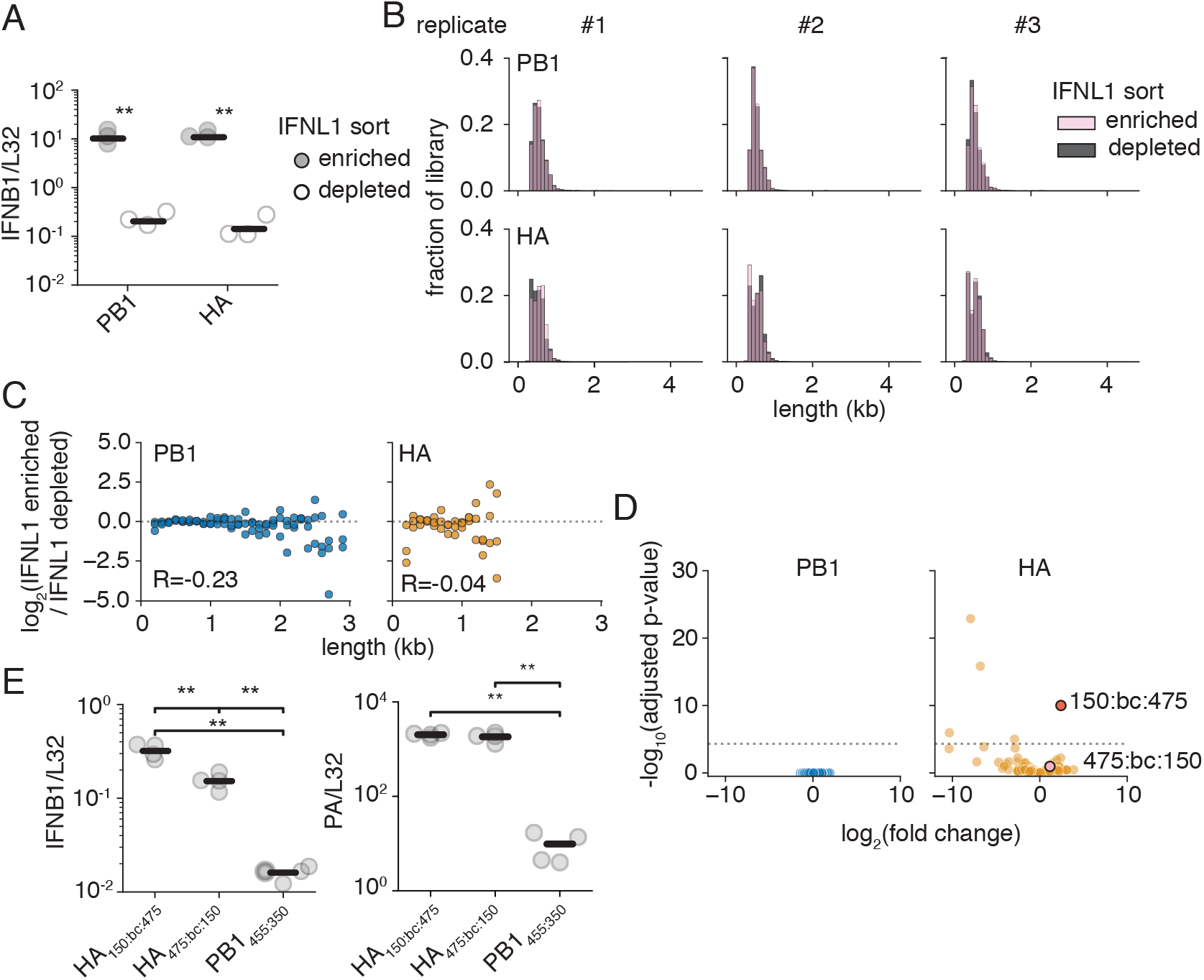
Deletions induce interferon in a length-independent manner. **(A)** An A549 type III interferon reporter line was infected at an MOI of 0.05 and sorted using magnetic-activated cell sorting 14h post infection. qPCR against the type I interferon, *IFNB1*, as corrected by the housekeeping gene *L32*. Asterisks indicate significantly increased *IFNB1* transcript, two-tailed t-test. n=3. **(B)** Size distributions of libraries from the sorting in **(A). (C)** Interferon induction and length are not correlated. Each point represents a 100nt bin. The fraction of variants falling within each bin was compared between interferon-enriched, and -depleted selections. Points above the dotted line are co-enriched with interferon. Points were only shown if represented in all three libraries under both conditions. R is the Spearman correlation coefficient. n=3. **(D)** Enrichment or depletion of individual variants within libraries. Each individual point represents a distinct 5’ and 3’ junction with <100 average observations. An adjusted p-value threshold of 0.05 is shown by the dotted line. Variants chosen for further analysis highlighted in red and pink. **(E)** Variants of identical length can exhibit different levels of interferon induction. (left)qPCR measuring levels of *IFNB1* normalized by *L32* in A549 cells infected at an MOI of 0.5 at 14h post-infection. (right) qPCR measuring levels of *PA* to show similar levels of infection between HA variants. Asterisks represent significantly different values in all pairwise comparisons, two-tailed t-test. n=4. For panels **(A)**, and **(D)**, individual points are individual measurements, lines are mean values. For panels **(A)**, and **(D)** siginficance was calculated using Benjamini-Hochberg correction at an FDR of 0.05. **Figure 5–Figure supplement 1**. Validation of sort in ***Figure 5***A. **Figure 5–source data 1**. full differential variant expression data are in `DESeq2library.tsv`.

Comparing length distributions of our library in interferon-enriched versus interferon-depleted populations, we notice no obvious shifts as we did in our other selections (***Figure 5***B). To explore in greater depth, we looked for evidence for enrichment with interferon as a function of length to look for more subtle shifts that may drive interferon expression (***Figure 5***C). We see no clear relationship between size and interferon state, with no strong enrichment at any size, at Spearman correlation coefficients of -0.23 and -0.04, for PB1 and HA, respectively. Overall, we fail to find significant evidence that length is a determinant of interferon production.

To probe whether we captured any variation in interferon induction within our libraries, we explored how individual junctions rather than binned sizes behaved in our interferon enrichment sequencing (***Figure 5***D). Using DESeq2 to analyze our data, there was little detectable variation between interferon-enriched and -depleted datasets in our PB1 library (***Love et al., 2014***). However, we find slightly more variation within our HA library, and identify a single variant with statistically significant enrichment in our interferon-producing cells. This variant consists of a junction lying 150nt from the 5’ end of the HA vRNA, then our 57nt adapter, and lastly a downstream junction lying 475nt from the 3’ end of the HA vRNA, for a total length of 682nt (HA_150:bc:475_). Curiously, the mirror-image deletion, HA_475:bc:150_ exhibits mild, not statistically significant, enrichment.

This finding provided us with an opportunity to demonstrate that length is not the sole contributor to whether any given deletion is stimulatory or not. We cloned both HA_150:bc:475_ and HA_475:bc:150_, and grew each independently using complementing cells. Thereafter, we tested the capacity of viral populations bearing these mutations to stimulate interferon (***Figure 5***E). As predicted by our more comprehensive analysis,HA_150:bc:475_ induces approximately 2-fold higher transcription of *IFNB1* than HA_475:bc:150_, demonstrating that features other than length must contribute to the variation in interferon induction.

### Interferon induction by defective segments is sensitive to genome replication

Critically, if our differentially-stimulatory variants exhibit different affinities for RIG-I or engage in aberrant behavior that indirectly impacts RIG-I signaling, then we might expect that these behaviors would be exacerbated by genome replication. While deletions in the polymerase segments produce defective species that, on their own, cannot engage in genome replication due to a requirement for *de novo* polymerase synthesis, deletions in non-polymerase segments would not be subject to such restrictions (***Vreede et al., 2004***). Therefore, we must ask, is the immune stimulation in our HA deletions contingent upon their ability to engage in genome replication? Interestingly, both HA_150:bc:475_ and HA_475:bc:150_ exhibit induction of *IFNB1* above that of our less stimulatory PB1 variant, PB1_445:bc:350_, suggesting that both act as efficient triggers of innate immunity, even if their potency differs.

To determine whether genome replication may play a role in this process, we used the nucleoside analog, ribavirin, to suppress genome replication (***Scholtissek, 1976***; ***Vanderlinden et al., 2016***). While prior studies have used the translation inhibitor cycloheximide, cycloheximide enhances interferon signaling to even pure innate immune agonists, thus complicating conclusions using this inhibitor (***Tan et al., 1970***; ***Raj and Pitha, 1983***; ***Ringold et al., 1984***). To confirm that we are investigating the effects of genome replication alone, and not pleiotropic impacts on cellular pathways, we compare the impacts of ribavirin treatment concurrently on infections with our less stimulatory PB1 deletion, PB1_445:350_, which is unable to replicate, and infections with HA_475:bc:150_. Upon ribavirin treatment, HA_475:bc:150_ exhibits a massive drop in viral transcription of the PA gene, consistent with reduced template vRNA molecules for mRNA synthesis (***Figure 6***A). PB1_445:350_ exhibits no drop as any mRNA transcribed in this virus must be from the initiating viral genome. With matched transcription levels between our deletions in HA and PB1, we now see a decrease in interferon induction for HA_475:bc:150_ but none for PB1_445:350_, demonstrating that genome replication is a key determinant of interferon induction even for defective influenza species.

**Figure 6.**
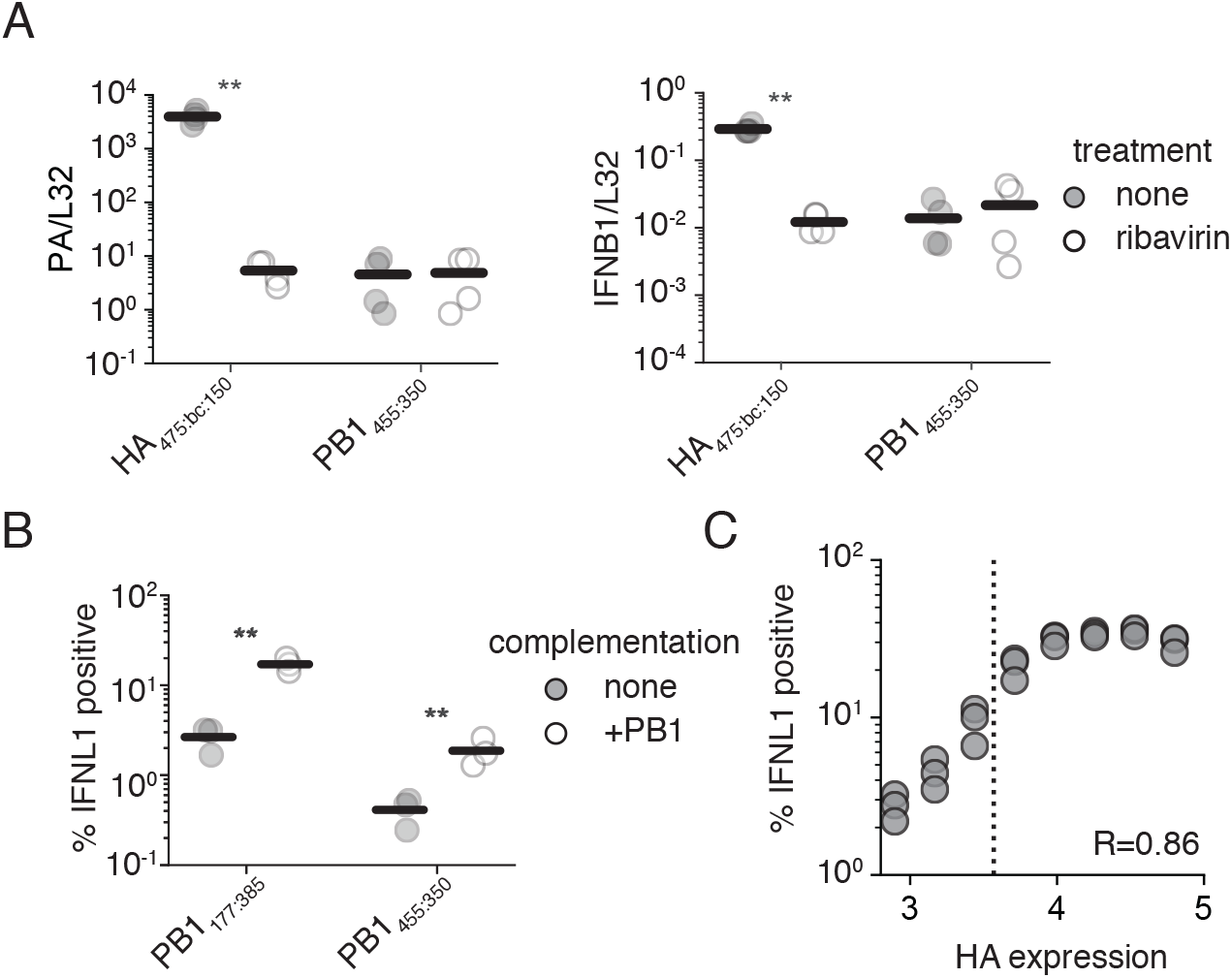
Genome replication is a key component of interferon induction. **(A)** (left) qPCR against *PA*, as corrected by the housekeeping gene *L32*, was used to validate the suppression of viral replication by the antiviral nucleoside ribavirin. (right) qPCR against *IFNB1* as corrected by the housekeeping gene *L32* demonstrates the suppression of interferon induction by a deletion capable of genome replication. A549 cells were pretreated with 200 μM ribavirin for 2h and then infected at an MOI of 0.5. RNA was harvested for analysis after 14h of infection. Astrisks indicate strains wherein signal was significantly impacted by treatment. Two-tailed t-test. n=4. **(B)** Complementation of PB1 defective species by PB1 expression *in trans* increases interferon induction. The fraction of cells in a population expressing a type III interferon reporter when infected with the indicated influenza variant at an MOI of 0.5 for 13h. Gating schema shown in ***Figure 6–Figure Supplement 1*. (C)** The increase in interferon-induction observed in **(B)** is linked to influenza replication. Complemented cells were infected as in **(B)** with an MOI of 0.5 of PB1_177:385_, stained for the viral protein HA and analyzed after 13h. Points represent binned values of HA expression, each point represents the right-most edge of a bin. The dotted line is the 99^th^ percentile of PB1_177:385_ HA staining without complementation. R represents the Spearman correlation coefficient. n=3. Gating schema shown in ***Figure 6–Figure Supplement 2*** For panels **(A)**, and **(B)**, individual points are individual measurements, lines are mean values. For panels **(A)**, and **(B)** siginficance was calculated using Benjamini-Hochberg correction at an FDR of 0.05. **Figure 6–Figure supplement 1**. Gating for ***Figure 6***B. **Figure 6–Figure supplement 2**. Gating for ***Figure 6***C.

This leads to an interesting question; would PB1_445:350_, noted to have approximately the same capacity to stimulate interferon as wild-type influenza, actually be potently stimulatory if it was capable of replicating its genome? To answer this question, we integrated a PB1 expression cassette into our type III interferon reporter line. This allowed us to measure the effects of complementation on interferon induction by flow cytometry, permitting a more refined measurement delineating whether complementation not only drives increased interferon induction, but also whether it fundamentally does so by increasing the number of cells that successfully detect influenza.

As expected from our model derived from ribavirin treatment, complementation of either PB1 deletion, PB1_445:350_ or PB1_177:385_, with PB1 expression *in trans* greatly increases the fraction of cells participating in the production of interferon, although each variant retains its relative level of stimulation to one-another (***Figure 6***B). This supports our conclusions that relative interferon induction is not only contingent on the amount, but also the sequence, of the aberrant segment. Combined with our prior observations, we can see that even those deletions that drive modest interferon induction, such as PB1_445:350_, could be quite dangerous to the virus were they to occur in non-polymerase segments.

Following from these data, we wanted to conclusively demonstrate that enhanced replication drives a higher probability an infected cell detects viral infection, and that our complementation experiments weren’t simply leading to additional rounds of infection. Using the highly stimulatory PB1_177:385_, we repeated the infection of complemented interferon reporter cells and concurrently stained for the viral protein HA. As transcription and replication are fundamentally linked in influenza viruses, increased protein production is highly correlated with genome replication (***Bean and Simpson, 1973***; ***Velthuis et al., 2018a***). Therefore, we may use HA staining as a rough proxy for the level of replication within an infected cell. In doing so, we see a positive relationship between the level of HA staining of infected cells and the probability that those cells produce interferon, with a Spearman coefficient of 0.89. This demonstrates that our results were not merely driven by an increased fraction of infected cells, but rather by an increased probability of response in cells with active viral replication (***Figure 6***C). Indeed, at higher levels of HA staining we find that PB1_177:385_, can induce interferon in as many as 35% of infected cells, rivaling prior measurements of interferon expression in viral infections lacking the critical interferon antagonist NS1 (***Killip et al., 2017***).

### Coinfection increases interferon induction by defective particles

Seeing that complementation appears to impact innate immune detection of defective polymerase species, we wondered whether this observation could be extended to coinfections, wherein intact polymerase genes are delivered by other viral particles, rather than an integrated expression cassette. In a coinfection we would expect some level of competition between our defective segment and its full-length counterpart, potentially reducing levels of RIG-I ligand. To explore whether coinfection reproduces data consistent with our complementation experiments, we turned to a pre-existing pseudovirus system in which the coding sequence of influenza HA has been replaced with mCherry. By infecting reporter cells with this pseudovirus alongside either wild-type influenza, or PB1_177:385_, we can identify coinfections as those that stain positive for both HA and mCherry (***Martínez-Sobrido et al., 2009***; ***Russell et al., 2018***). Curiously, our pseudovirus alone appears to induce much higher levels of interferon than wild-type influenza, potentially due to the presence of non-native sequence. Regardless, when present in a coinfection with PB1_177:385_ a greater fraction of cells express interferon than when infected with either virus alone (***Figure 7***A). A similar effect is not seen for wild-type influenza, demonstrating the specificity of this phenomenon to defective influenza species. Thus complementation during coinfection can drive higher levels of interferon induction by deletions in polymerase segments.

**Figure 7.**
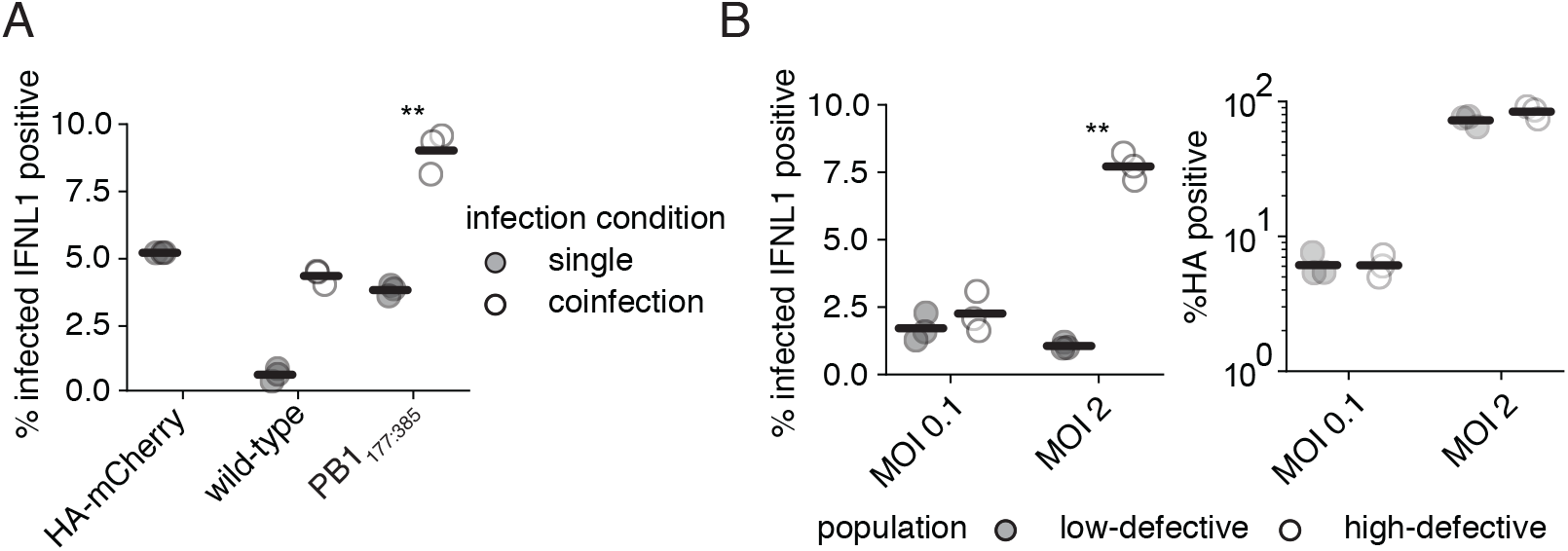
Coinfection increases interferon induction by defective particles. **(A)** Coinfection with a complementing virus increases interferon induction by PB1_177:385_. Type III interferon reporter A549 cells were infected with PB1_177:385_ or wild-type influenza at an MOI of 0.1, and HA_mCherry_ pseudovirus at an MOI of 0.5. Interferon expression was measured in HA-stained cells at 13h post infection. Cells from single infections were called as infected if they stained positive for HA or expressed mCherry. Coinfections required co-staining for both HA and mCherry. Asterisks indicate conditions under which coinfection led to a significant increase in interferon from both HA_mCherry_ and the indicated variant alone, one-tailed t-test. Gating schema shown in ***Figure 7–Figure Supplement 1*** for HA and mCherry, and ***Figure 7–Figure Supplement 2*** for interferon. **(B)** Populations replete in defective particles exhibit much higher interferon induction under high MOI infections where coinfection is common. (left) The fraction of HA-positive cells expressing interferon. (right) The total fraction of HA-positive cells. Type III interferon reporter A549 cells were infected with a low-defective or high-defective population as described in ***Figure 4–Figure Supplement 1***. Cells were then infected at a genome-normalized MOI as calculated from the low-defective population and stained for HA and analyzed for interferon reporter expression after 13h. Gating schema shown in ***Figure 7–Figure Supplement 3***. For panels **(A)**, and **(B)**, individual points are individual measurements, lines are mean values. For panels **(A)**, and **(B)** siginficance was calculated using Benjamini-Hochberg correction at an FDR of 0.05. **Figure 7–Figure supplement 1**. Gating for ***Figure 7***A for HA and mCherry. **Figure 7–Figure supplement 2**. Gating for ***Figure 7***A interferon. **Figure 7–Figure supplement 3**. Gating for ***Figure 7***C.

Following the logic that complementation from co-infecting viruses can lead to increased detection of defective influenza particles, we might expect that deletions in the polymerase segments drive a proportionately greater innate immune response at higher infectious doses where coinfection is more common. To see if this assumption is true we infected our interferon reporter line at an MOI of either 0.1 or 2 of a low-defective population, and equivalent genome copies of a high-defective population, and measured the fraction of HA-positive cells that were expressing interferon lambda (***Figure 7***B). With a low-defective particle population, the average rate at which infected cells produce interferon remains largely unchanged between these two MOIs. However, with a high defective population, there is a massive increase in the fraction of interferon-producing cells at a higher MOI. This was not due to a difference in infection rate, as the fraction of HA-expressing cells in each condition between the low and the high defective content viral populations was similar. This suggests that interferon induction by the vast majority of defective species is a nonlinear process. Their relative importance is dependent upon not only their frequency, but also the rate of complementation as driven by coinfection by viruses bearing full-length segments. Critically, these processes modulate interferon induction, but genome replication is not absolutely required for detection of defective species as demonstrated by stimulation from PB1_177:385_ alone. Thus apparent contradictions in the field are likely real, meaningful, differences, attributable to choice of MOI, the fraction of a given population that is defective, and the sensitivity of the method used to detect interferon production rather than any actual discrepancies in the underlying biological pathways (***Österlund et al., 2012***; ***Rehwinkel et al., 2010***; ***Liu et al., 2019***; ***Fay et al., 2020***; ***Killip et al., 2014***).

## Discussion

We have used a library-based approach to comprehensively explore the relationship between length and replication speed, packaging, and interferon induction in influenza A virus for the polymerase segment, PB1, and the non-polymerase segment, HA. Additionally, we describe how genome replication can contribute to interferon induction by defective species, and demonstrate that both the population composition and the rate of coinfection influence the probability with which these viruses are detected by cell intrinsic innate immunity (***Figure 8***).

**Figure 8.**
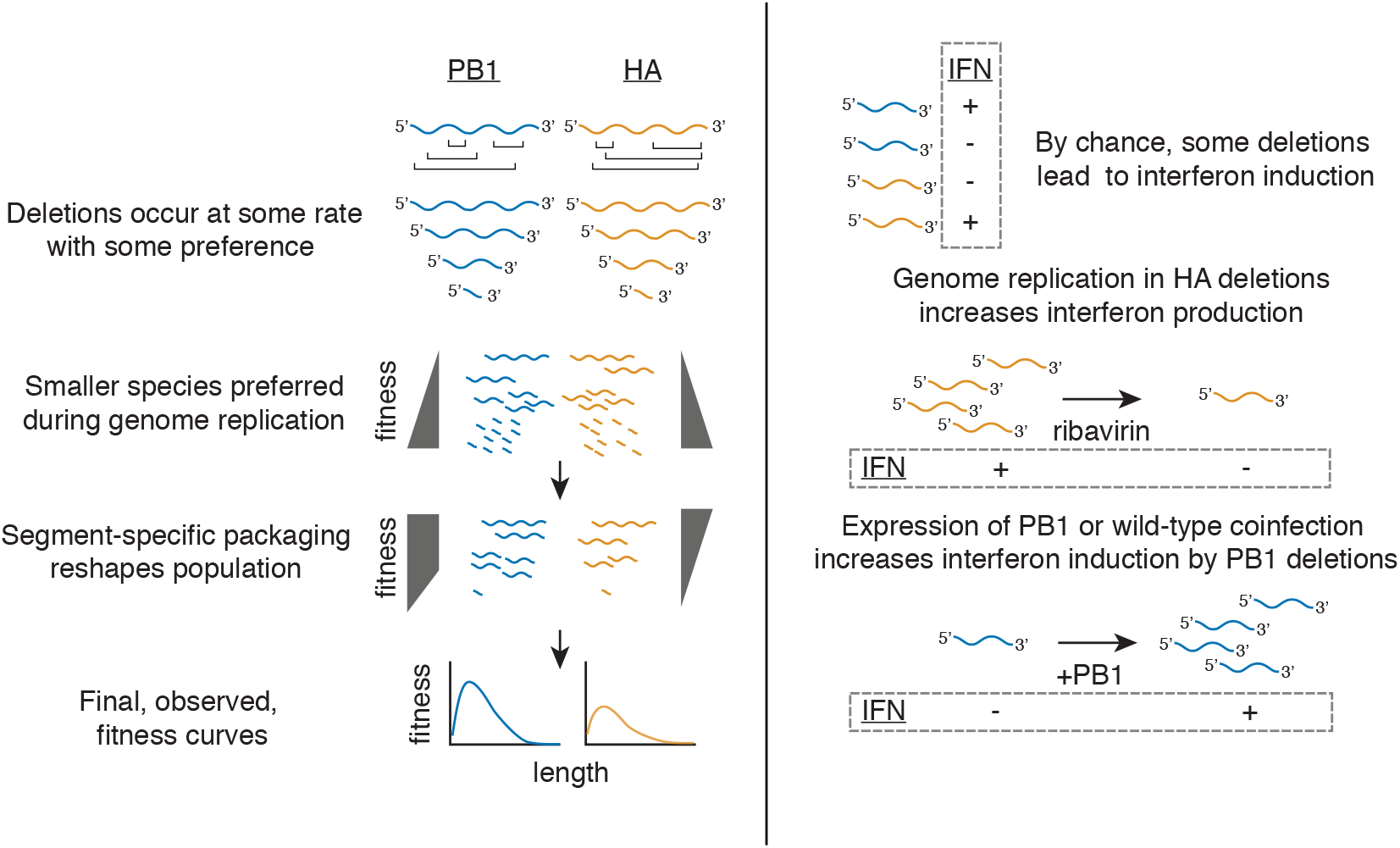
Summary of major conclusions. After deletions form, they are subject to selection during genome replication, which favors smaller products. Packaging in the PB1 segment excludes very small products, but does not discriminate after ∼400nt of sequence. Packaging in the HA segment exhibits a length dependence throughout, favoring larger products. Combining pressures produces a curve with a deletion fitness maximum of ∼400 nt, but with a lower overall preference for HA deletions than PB1 deletions. Interferon induction by deletions is a length-independent process. Genome replication modulates inteferon induction by deletions, either occuring in single infections for non-polymerase deletions, or during coinfection for polymerase deletions.

Interestingly, during replication and viral packaging our artificial libraries converge on a near-identical length as natural defective populations (***Pelz et al., 2021***). This suggests that the contrasting pressures of replication speed and packaging are sufficient to shape defective influenza populations to their final, observed, lengths, from potentially disparate initial distributions. Therefore, we cannot truly infer the rates or locations of *de novo* deletion formation from a fully-matured viral population. For instance, the existence of rare, small, deletions that appear with, and likely hitchhike on, larger deletions is consistent with an understanding that *de novo* deletion formation may bear little resemblance to final viral populations (***Sivasubramanian and Nayak, 1983***; ***Saira et al., 2013***).

The divergence in packaging between PB1 and HA also underscores the capacity of this step in the viral lifecycle to shape the distribution of defective viral segments. A recent report described a similar finding; that HA readily accumulates deletions within cells, but that these deletions are generally lost during packaging (***Alnaji et al., 2021***). Our study, which provides length variation above that found in natural populations, provides an explanation in the form of divergent requirements in the amount of sequence necessary for efficient packaging. This suggests that diffuse contacts are much more important for HA than PB1, and that differences between these two segments are due to specific, discriminatory, mechanisms, rather than chance biophysical properties such as the marginal difference in length between HA and PB1.

In contrast to replication speed and packaging, we were surprised to find that interferon induction by influenza A virus defective particles is a length-independent process. This finding is consistent with similar work on *Paramyxoviridae*, which generate short, copy-back defective genomes. Work on these viruses found that it was not the length of these genomes, nor their double-strandedness, that leads to their detection, but rather changes in their RNA secondary structure (***Xu et al., 2015***). It seems likely that deletions in influenza likewise perturb otherwise non-stimulatory sequence composition, and, by chance, these perturbations can generate potent RIG-I ligands. The finding that small defective genomes preferentially associate with RIG-I is consistent with this hypothesis based on the following logic: 1) sequence perturbation away from the wild-type concensus will generally be purged from the population, or at least will not rise to high levels, as it is likely to result in a loss of fitness, 2) defective genomes, while replication-incompetent, nevertheless can rise to high levels in the population owing to their ability to act as parasites and out-compete full-length genomes during genome replication, and therefore 3) sequence perturbation that drives interferon stimulation is most likely to be found in defective genomes. Thus, length would be observed to correlate well with interferon induction in natural populations, even if length was not the underlying causal factor. This does not preclude the role of mvRNAs <125nt in interferon induction, and these incredibly small RNA species likely contribute alongside defective genomes to the observed interferon response (***Velthuis et al., 2018b***; ***Nilsson-Payant et al., 2021***).

Lastly, we find that replication contributes to, but is not necessary for, the innate immune response to defective influenza particles. This is in slight contrast to several reports that genome replication fails to modulate innate immune detection of these populations. This seeming discrepancy is easily attributable to two experimental decisions: 1) the use of cycloheximide as an inhibitor, which is known to increase the transcription of interferons when cells are challenged with the viral mimetic poly I:C, thus making it difficult to understand whether increases, or lack of decrease, in response to influenza is due to perturbations in the viral lifecycle or alteration of cellular signaling pathways (***Killip et al., 2014***; ***Tan et al., 1970***; ***Raj and Pitha, 1983***; ***Ringold et al., 1984***) and 2) some of these studies use MOIs of high-defective populations as high as 50, which may represent particle doses of 5,000-50,000/cell (***Liu et al., 2019***). At such high doses it is quite likely that additional ligands presented by genome replication would only play, at most, a modest role in interferon induction.

The finding that genome replication modulates interferon induction in turn leads us to reconsider the observed distribution of defective species across the viral genome, largely consisting of deletions in the polymerase segments, in a new light. Under conditions early in infection, where coinfection may be more limited, occasional packaging of deletions in the polymerase segment may be tolerated as the likelihood these deletions induce interferon remains low. However, should the virus package non-polymerase deletions, these viruses would exhibit an increased probability of interferon induction even under conditions where coinfection is rare. We therefore postulate that, rather than defective particles preferentially consisting of deletions in the polymerase segments due to some selection for their inclusion, there may simply be a strong selection against the packaging of deletions in non-polymerase segments.

Overall our study addresses several critical gaps in our understanding of influenza A virus defective particles. Our library-based approach approach was quite successful at generating relatively complete models of how genome replication and packaging shape defective influenza populations. Although we were only able to make a few inferences regarding the precise nature of deletions that drive interferon induction, namely that this phenomenon is length-independent and impacted by genome replication, our study was not designed to probe sequence-dependent phenomena as well as length-dependent owing to the introduction of barcodes and other artificial sequences which themselves likely impacted RNA structure. Future work with a modified library design will be required for further exploration of those features with an ultimate goal of being able to perform *in silico* prediction of stimulatory behavior. Finally, beyond this work on influenza A virus, we note that our library design is also broadly extensible to explore any length-dependent phenomena wherein the lengths are compatible with PCR amplification, and hope that providing this method will serve as a resource to the broader community.

## Materials and Methods

### Cell lines and viruses

The following cell lines were used in this study: HEK293T (ATCC CRL-3216), MDCK-SIAT1 (variant of the Madin Darby canine kidney cell line overexpressing SIAT1 (Sigma-Aldrich 05071502)) and A549 (human lung epithelial carcinoma cell line, ATCC CCL-185). Variants of MDCK-SIAT1 overexpressing influenza HA and PB1 were generated using a lentiviral vector as previously described (***Doud and Bloom, 2016***; ***Bloom et al., 2010***; ***O’Connell et al., 2010***) A549 type I and type III interferon reporter lines were previously described (***Russell et al., 2019***). A549 cells overexpressing PB1 were generated using lentiviral transduction using the same vector for MDCK-SIAT1. Cell lines were tested for mycoplasma using the LookOut Mycoplasma PCR Detection Kit (Sigma-Aldrich) using JumpStart Taq DNA Polymerase (Sigma-Aldrich). A549 cells used in all sequencing experiments had their identity confirmed via STR profiling by ATCC. All cell lines were maintained in D10 media (DMEM supplemented with 10% heat-inactivated fetal bovine serum and 2 mM L-Glutamine) in a 37°C incubator with 5% CO_2_.

Wild type A/WSN/1933 (H1N1) virus was created by reverse genetics using plasmids pHW181-PB2, pHW182-PB1, pHW183-PA, pHW184-HA, pHW185-NP, pHW186-NA, pHW187-M, pHW188-NS (***Hoffmann et al., 2000***). Genomic sequence of this virus provided in `WSNgenome.fa`. HEK293T to MDCK-SIAT1 cells were seeded in an 8:1 coculture and transfected using BioT (Bioland Scientific, LLC) 24h later with equimolar reverse genetics plasmids. 24h post transfection, D10 media was changed to Influenza Growth Medium (IGM, Opti-MEM supplemented with 0.04% bovine serum albumin fraction V, 100 μg/ml of CaCl_2_, and 0.01% heat-inactivated fetal bovine serum). 48 post-transfection, viral supernatant was collected, centrifuged at 300g for 4 minutes to remove cellular debris, and aliquoted into cryovials to be stored at -80°C. Thawed aliquots were titered by TCID50 on MDCK-SIAT1 cells and calculated using the Reed and Muench formula or titered by qPCR against the viral segment HA (***REED and MUENCH, 1938***). To create wild type viral stocks with a low defective content, MDCK-SIAT1 cells were infected at an MOI of 0.01 and harvested 30 hours post-infection. To generate intermediate defective populations, MDCK-SIAT1 cells were serially infected at an MOI of 0.01 twice, harvested at 72h each passage. To generate high-defective populations, these passages were instead performed at an MOI of 1.

Indicated mutations were introduced by inverse-PCR followed by single-molecule Gibson ligation and confirmed by Sanger sequencing. To generate viruses bearing large deletions in PB1 or HA, HEK293T cells were co-transfected with all reverse-genetics plasmids encoding A/WSN/1933 except for either HA, or PB1, the indicated variant, and a construct expressing either HA or PB1 under control of the a constitutive promoter in the pHAGE vector as used to transduce MDCK-SIAT1 cells. MDCK-SIAT1 cells in co-culture either constitutively expressed HA or PB1. Viruses were titered on complementing MDCK-SIAT1 cells, or by qPCR against the viral segment PA. For the barcoded HA deletions, HA_150:bc:475_ and HA_475:bc:150_, 5’ AGCTCCTATTAG 3’, and 5’ TATAGTCCCAAT 3’, were used instead of random sequence.

### Library cloning and initial sequencing

Up (amplifying the 5’ end of the gene) and Down primer sets were used in individual PCR reactions alongside the common primers 5’ GGTAACTGTCAGACCAAGTTTACTC 3’ (Up) or 5’ GAGTAAACTTG-GTCTGACAGTTACC 3’ (Down) using Q5 Hot Start High-Fidelity 2x Master Mix (New England Biolabs, M0494). Primers were designed as described in https://github.com/a5russell/Defective_Library_Mendes_Russell, using annealing temperature calculations as originally described in (***Breslauer et al., 1986***). Each reaction was performed using 2.5 ng of template DNA consisting of a pHH21 vector encoding either HA or PB1 with the ATG start codon mutated to GTA (***Neumann et al., 1999***). Reaction conditions consisted of a 55°C annealing temperature, a 2 minute extension, and 11 total cycles of amplification. Up and Down amplicons were independently pooled and subjected to DpnI digest at 37°C for 1 hour. Products were then purified using SPRI beads from the ProNex Size-Selective Purification System (Promega, NG2001) at a 1x bead volume following manufacturer’s instructions.

Down amplicons were then subjected to a second round of PCR using the same common PCR primer and an additional primer consisting of an artificial adapter containing a 12nt random nucleotide sequence, 5’ GCCATCGCCTACACGACGCTTCNNNNNNNNNNNNCGTTGACCACTGCTTGCGATGAT 3’. This primer was acquired from integrated DNA technologies, with a hand-mixed randomer. Conditions for this amplification consisted of 25ng of template DNA subjected to 8 cycles of amplification at a 50°C annealing temperature and 2 minute extension. This amplicon was then purified once more using a 1x bead volume.

25 ng of each amplicon pool, Up and Down with adapter, were combined in a Gibson reaction using NEBuilder HiFi DNA Assembly Master mix (New England Biolabs, E2621) using manufacturer’s protocols save for a 1 hour incubation step rather than 15 minutes. The resultant reaction was then cleaned using 1x bead volumes, and electroporated into electrocompetent Stbl3 cells. Plates were counted, scraped, and plasmids prepared for each step. This whole process, from PCR onwards, was performed 3 independent times for each library.

To sequence junction regions associated with each barcode, 1μg of each library was digested with BtgZI overnight and released products were purified by gel electrophoresis followed by gel extraction. Bands were then subjected to ligation with a DNA splint containing an NNNN region to facilitate ligation to the released overhang. This splint was generated by annealing 5’ AAGCAGTG-GTATCAACGCAGAGTACAT 3’ to 5’ NNNNATGTACTCTGCGTTGATACCACTGCTT 3’. Ligation to each splint was performed overnight at 16°C using T4 DNA ligase (New England Biolabs).

Ligation reactions were used directly as templates for PCR to infer Up junctions and Down junctions, separately, using the common primer 5’ CGGGGTACGATGAGACACCATATTGGTCTCAAGCAGTG-GTATCAACGCAG 3’, and 5’ GAAGCAGAAGACGGCATACGAGATAGGCCATCGCCTACACGACGCT 3’ (Up) or 5’ GAAGCAGAAGACGGCATACGAGATAGATCATCGCAAGCAGTGGTCA 3’ (Down) with the following reaction conditions: 60°C annealing temperature, 15s extension, for 12 cycles. A second round of PCR was then used to add partial Illumina adapters (5’ TCGTCGGCAGCGTCAGATGTGTATAAGA-GACAGCGGGGTACGATGAGACACCA 3’ / 5’ GTCTCGTGGGCTCGGAGATGTGTATAAGAGACAGGAAGCA-GAAGACGGCATACGAGAT 3’, with a 55°C annealing temperature, 15s extension, and 6 cycles. Samples were then barcoded using Illumina UD indices with a 62°C annealing temperature, 20s extension, and 10 cycles. HA and PB1 libraries were pooled in indexing as they could be disambiguated during sequencing. Final samples were bead cleaned with a 3x bead volume and sizes confirmed by tapestation prior to sequencing.

### Barcoded library selections

To analyze rates of genome replication, in an 8:1 coculture of HEK293T and MDCK-SIAT1 cells, each individual deletion/duplication library was cotransfected with plasmids that individually encode the mRNA for PB2, PB1, PA, and NP in an equimolar ratio using BioT according to the manufacturer’s protocol. An orthogonal dataset was generated with transfections consisting of a library, PB1, PA, and NP but not PB2. Transfections were then allowed to proceed for 24 hours and RNA was purified using the Monarch Total RNA Miniprep Kit from New England Biolabs (T2010). To analyze viral packaging, after 24h media was changed from D10 to IGM and cells were infected with a low-defective wild-type WSN population at an MOI of 0.25, and infection was allowed to progress for 72h. Virus-containing supernatant was clarified by centrifugation at 300g for 4 minutes, and RNA was purified from 100 μl of supernatant using the Monarch Total RNA Miniprep Kit with 600 μl of lysis buffer.

To associate barcodes with the induction of interferon, 21 million interferon-lambda reporter cells were seeded across 3 15cm plates. 24 hours later, the media was changed to IGM and cells were infected with library supernatant at an MOI of 0.05. 14 hours post-infection, cells were separated using the MACSelect LNGFR System (Miltenyi Biotec). The supernatant of the samples were aspirated before being trypsinized for 10 minutes at 37°C and quenched with D10 media. Samples were centrifuged for 4 minutes at 300g, had their supernatant removed, and were resuspended in 320 μL of 1x phosphate buffered saline (PBS). 80 μL of MACSelect LNGFR MicroBeads were mixed into the samples thoroughly and then placed in a 4°C fridge for 15 minutes. Samples were then applied to LS columns (Miltenyi Biotec, 130-042-401) according to manufacturer’s instructions. In addition to our enriched population, five million interferon-depleted cells from each sample were retained for analysis. RNA was thereafter purified from both sets of samples using the Monarch Total RNA Miniprep Kit from New England Biolabs.

### Barcoded library sequencing

All RNA samples (minigenome +PB2, minigenome -PB2, viral supernatant, and interferon-enriched and -depleted sorts) were converted to cDNA using the universal vRNA primers of Hoffmann *et al*. and the SuperScript III First-Strand Synthesis SuperMix kit (Invitrogen, 18080400) according to the manufacturer’s protocol (***Hoffmann et al., 2001***). Before converting the interferon sorted RNA samples to cDNA, a qPCR against the HA segment or PB1 segment was performed to equilibrate the number of RNA molecules to be reverse transcribed between each enriched and depleted biological sample. Primers for this qPCR were, for PB1, 5’ CCGTCTGAGCTCTTCAATGG 3’/5’ GATGCACGAATTGATTTCGAATCTG 3’, and for HA, 5’ GCAAAACTACTGGTCCTGTTATATGC 3’/5’ GTGTCAACAGTGTCGGTTGAG 3’.

cDNA samples were amplified and partial i5 and i7 adapters added with the following, sequential, amplifications using Q5 Hot Start 2x Master Mix (New England Biolabs), each consisting of 10 cycles and a 55°C annealing temperature with the following primer pairs: reaction 1, 5’ TGTGCTCTTCCGGCCATCGCCTACACGACGCT-3’/5’ AAGCAGTGGTATCAACGCAGAGTACATATCATCG-CAAGCAGTGGTCA 3’; reaction 2, 5’ GAAGCAGAAGACGGCATACGAGATAGTGTGCTCTTCCGGCCATCGC 3’/5’ CGGGGTACGATGAGACACCATATTGGTCTCAAGCAGTGGTATCAACGCAG 3’; reaction 3, 5’ TCGTCG-GCAGCGTCAGATGTGTATAAGAGACAGCGGGGTACGATGAGACACCA 3’/5’ GTCTCGTGGGCTCGGAGAT-GTGTATAAGAGACAGGAAGCAGAAGACGGCATACGAGAT 3’. A final PCR was performed to index all samples individually with a 62°C annealing temperature and 20 second extension time for 7 cycles Samples were purified by gel electrophoresis and agarose gel extraction. Products were pooled and cleaned by a 3x volume bead clean-up.

### Sequencing of naturally-occurring deletions in influenza A virus

Interferon-beta A549 reporter cell lines were infected at an MOI of 0.1 and sorted using magnetic activated cell sorting as described in the previous section as for interferon-lambda reporters, save for MS columns were used instead of LS columns. RNA was purified using either RNeasy plus mini kit (Qiagen, 74134), for interferon-depleted samples, or the RNease plus micro kit (Qiagen, 74034) for interferon-enriched samples. As above, RNA was converted to cDNA using the universal vRNA primers of Hoffmann *et al*. and the SuperScript III First-Strand Synthesis SuperMix kit (Invitrogen, 18080400) according to the manufacturer’s protocol. Samples were normalized by qPCR against HA. PCR was then performed as per Hoffmann *et al*. for 16 cycles.

To generate cDNA from mRNA, normalized RNA samples were converted to cDNA using the SMART-Seq2 method and subjected to 14 cycles of amplification (***Picelli et al., 2013***). Both mRNA and vRNA samples were fragmented by tagmentation using the Nextera XT DNA library preparation kit from Illumina (FC-131), indexed, and sequenced.

### Computational analysis of sequencing data

The entire analysis pipeline is found at https://github.com/a5russell/Defective_Library_Mendes_Russell. To analyze barcodes, a custom Python script was used to parse sequencing files. Upstream and downstream junctions were assigned to barcodes separately and combined to generate a barcode-junction map. Likely due to PCR chimerism, highly abundant barcodes ended up assigning to multiple junctions, but these secondary assignments were rare. Therefore, an empirical threshold of 75% of reads must confidently assign to a single junction to assemble a barcode. A single mismatch was allowed within junction sequence for assembly. Thereafter, for each individual sample the number of times a barcode was identified within sequencing was analyzed without any further discrimination, no score values needed to be considered, and mismatched barcodes were discarded. This resulted in an assembly for the majority of observed barcodes, suggesting little benefit could be procured using a more complex analysis pipeline allowing for a greater hamming distance. As we used paired-end reads for all sequencing, barcodes or junctions were collapsed to a consensus sequence, if a mismatch was observed whichever read possessed the higher quality score at that position was presumed to be the correct nucleotide.

To analyze mRNA sequencing, reads were first trimmed using Trimmomatic providing the Nextera adapter sequences and the following variables: 2 seed mismatches, a palindrome clip threshold of 30, a simple clip threshold of 10, a minimum adapter length of 2, keep both reads, a lead of 20, a sliding window from 4 to 15, and a minimum retained length of 36 (***Bolger et al., 2014***). Reads were then mapped against a concatenated GRCh38 human genome assembly and the A/WSN/1933 genome using STAR with default settings (***Dobin et al., 2013***). HTseq-count was used to count occurrences of a given transcript within our read mapping (***Anders et al., 2015***). Output from HTseq-count was then used in DeSeq2 using a short R script, and results parsed using Python scripts.

To analyze vRNA sequencing, reads were also trimmed using Trimmomatic as above. However, next read1 and read2 were mapped separately using STAR against the A/WSN/1933 genome with a requirement to map without any gaps by enforcing mapping to a custom gtf file consisting of the full length of each vRNA. Mismatches were excluded with the command –outFilterMismatchNmax of 3, deletions were largely excluded by providing a deletion open penalty of 6, and deletion per base penalty of 6. Unmapped reads were retained, and were then used in a BLASTn search against a custom BLAST database consisting of just the A/WSN/1933 genome (***Altschul et al., 1990***). Input for this search used a percent identity of 90, a word size of 10, a gap open penalty of 5, extend penalty of 2, and an evalue cutoff of 0.000001. BLAST output was then parsed and reads were identified that mapped to the same segment, discontinuously. Such reads must map completely, with no “missing” bases, and must map with the same polarity, with no inversion. Repetitive elements, such as 2nt repeated at the 5’ and 3’ ends of the junction, which could, in theory, be assigned to either, were assigned to the 5’ end arbitrarily.

BLAST output was then used to initialize a new gtf file, using all data from all technical replicates for a given biological replicate. Therefore two gtf files were generated, one for each biological replicate. Unmapped reads were then mapped using this updated file using STAR with identical parameters as above. The resulting four files, two from the first mapping and two from the second, updated, mapping, were then combined and bam flags were fixed for appropriate, paired-end, analysis. Junctions were thereafter counted in the bamfile with a requirement that both reads mapped, neither is inconsistent with the deletion (ie if both map over the region a deletion is contained within, both show a deletion within the read), and that at least three bases are mapped on either side of the junction when considering the consensus produced by the paired-end reads.

Output was then parsed using custom Python scripts and Samtools (***Li et al., 2009***).

### qPCR

Code generating graphs in manuscript can be found at https://github.com/a5russell/Defective_Library_Mendes_Russell. Primers for all qPCR analyses listed in `qPCRPrimers.tsv`. For validation of natural diversity, purified RNA was used as described above. For all other qPCR analyses, the method of Shatzkes *et al*. was used (***Shatzkes et al., 2014***). In brief, adherent cells were incubated, after a PBS wash, with a gentle permeabilization buffer consisting of 10 mM Tris-HCl, 150 mM NaCl, 0.25% IGEPAL, and 1% DNaseI in DNase/RNase-free water and incubated at 37°C for 5-10 minutes. DNase and infectious virus were then inactivated with a 5 minute 85 degree Celsius incubation. For supernatant measurements, 10 μl of supernatant was incubated with 90 μl buffer instead.

cDNA from lysate was generated using the High Capacity First Strand Synthesis Kit (Applied Biosystems, 4368814) with random hexamers according to the manufacturer’s protocol using a final lysate concentration of 10% of the reaction volume. qPCR was thereafter performed using Lunascript Universal qPCR Master Mix (New England Biolabs, M3003) with manufacturer’s suggested reaction conditions. For pure plasmid controls, plasmids were used at the indicated concentrations instead of cDNA in qPCR reactions.

### Flow cytometry

Indicated cells were seeded 24 hours prior to infection, and, at the indicated time points, trypsinized and resuspended in PBS supplemented with 2% of heat-inactivated fetal bovine serum (FBS). For HA staining, cells were stained with 10 μg/ml of H17-L19, a mouse monoclonal antibody confirmed to bind to WSN HA in a prior study (***Doud et al., 2017***). Cells were refrigerated with the primary antibody for one hour at 4°C before being washed with PBS supplemented with 2% FBS and then stained with a goat anti-mouse IgG antibody conjugated to allophycocyanin and refrigerated for another hour at 4°C. Cells were washed with PBS supplemented with 2% FBS and fixed with 1% formaldehyde (BD Cytofix) for 30 minutes at room temperature. Cells were washed with PBS and thereafter run on a flow cytometer. Data processing consisted of first generating a debris gate in FlowJo prior to export to a tsv or csv and analysis using custom Python scripts.

## Supporting information

DESeq2.tsv

DESeq2library.tsv

libraryPrimers.tsv

qPCRPrimers.tsv

## Data availability

Pipeline can be found at https://github.com/a5russell/Defective_Library_Mendes_Russell. All sequencing data can be found under the BioProject accession PRJNA760790, and will be made accessible upon publication of this manuscript.

## Acknowledgments

We thank the members of the Russell laboratory, and Jacquelyn Braggin, for comments on the manuscript. This work was supported by the NIAID of the NIH under grant K22 AI141678 and by the Damon Runyon Cancer Research Foundation Dale F. Frey Award, DFS 36-19. The funders had no role in study design, data collection and analysis, decision to publish, or preparation of the manuscript.

---

**Figure 2–Figure supplement 1.**
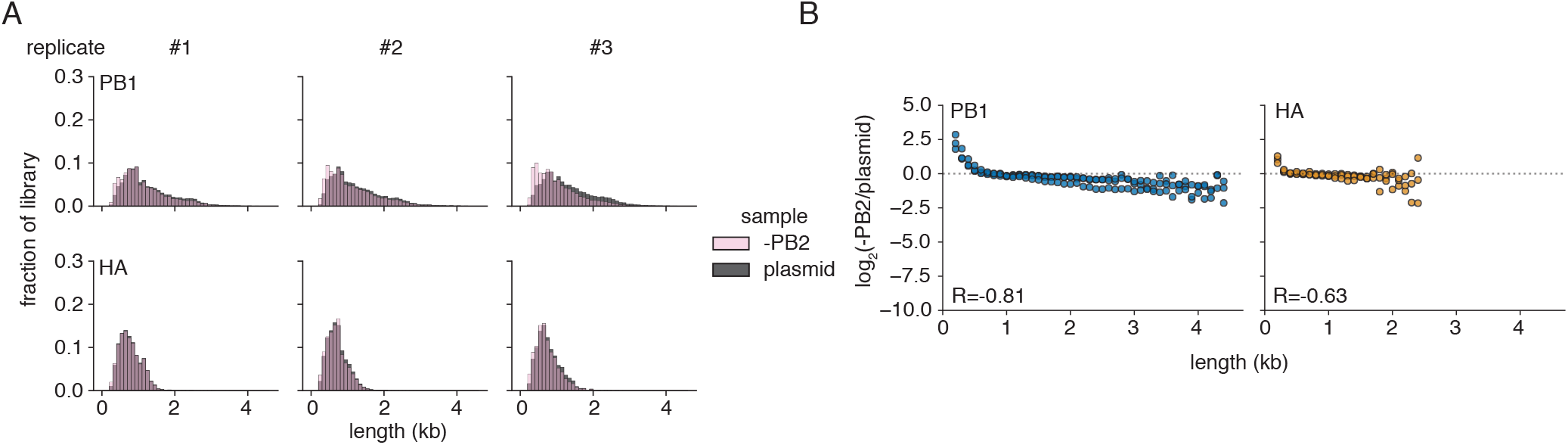
Biases present from polymerase I transcription alone. **(A)** Size distributions of original plasmid libraries and matched length distribution after transfection in the absence of the full viral genome replication machinery (-PB2) **(B)** Polymerase I transcription alone leads to a slight length bias in library composition. To analyze enrichment, or depletion, the fraction of variants falling within each 100nt bin was compared between replication-incompetent selection and the original plasmid library. Points above the dotted line represent lengths which were overrepresented in our polI transcribed library relative to a plasmid control. Points were only shown if represented in all three libraries under both conditions. R-value shown is the Spearman correlation coefficient. n=3.

**Figure 2–Figure supplement 2.**
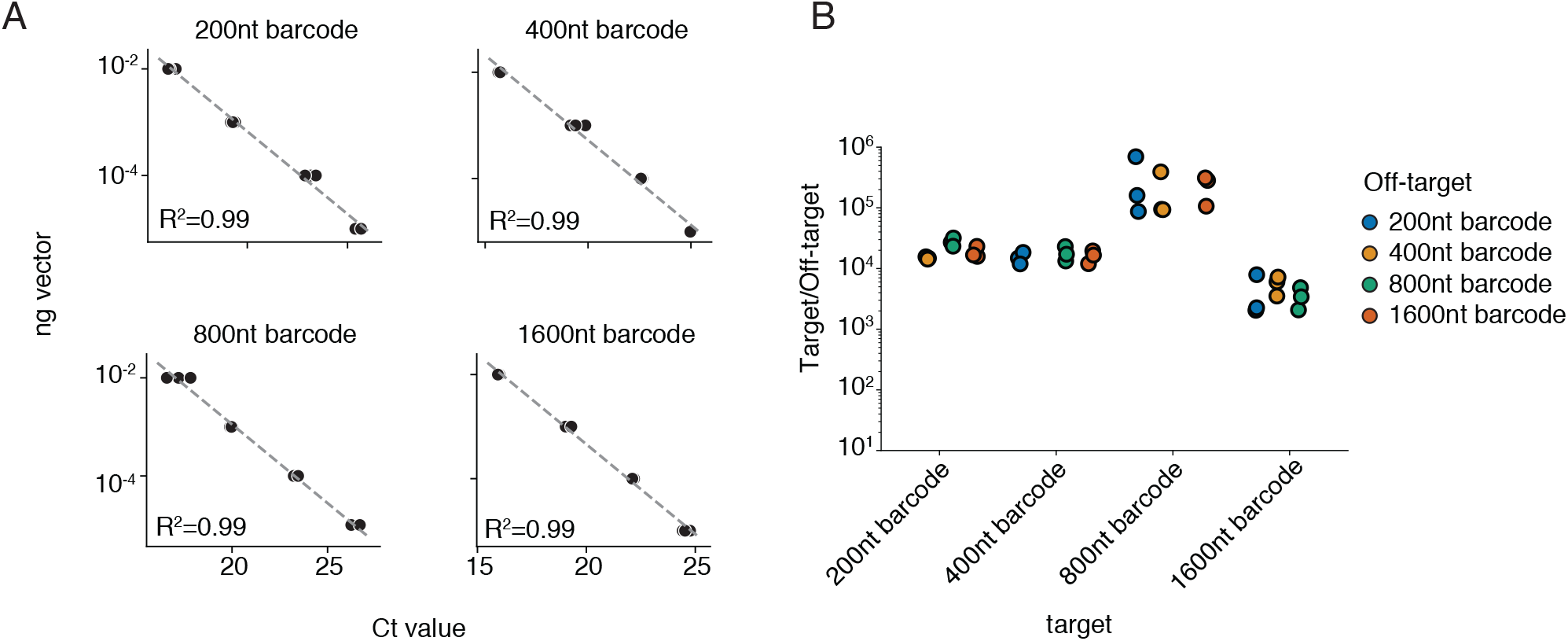
Validation of qPCR used in **Figure 2**C. **(A)** All four qPCR reactions exhibit linearity when tested against a plasmid control. R^2^ from linear regression against Ct verus the log-transformed plasmid concentration. Linear regression line shown. **(B)** Each qPCR is highly-specific to its cognate target, and discriminatory against non-cognate targets. Target versus non-target specificity was calculated as the relative signal of each qPCR reaction on the indicated off-target control when compared to a target control at 0.01 ng per qPCR reaction. n=3 for both panels.

**Figure 3–Figure supplement 1.**
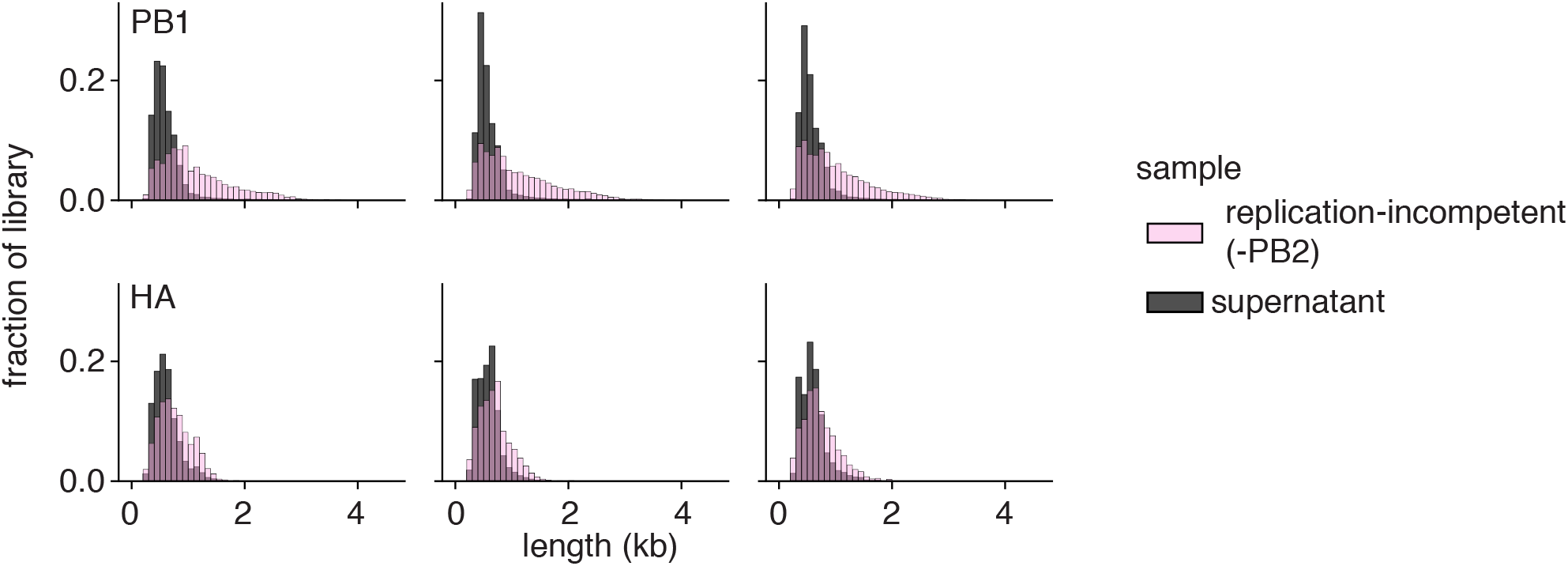
Size distributions used to calculate enrichment in **Figure 3**D.

**Figure 4–Figure supplement 1.**
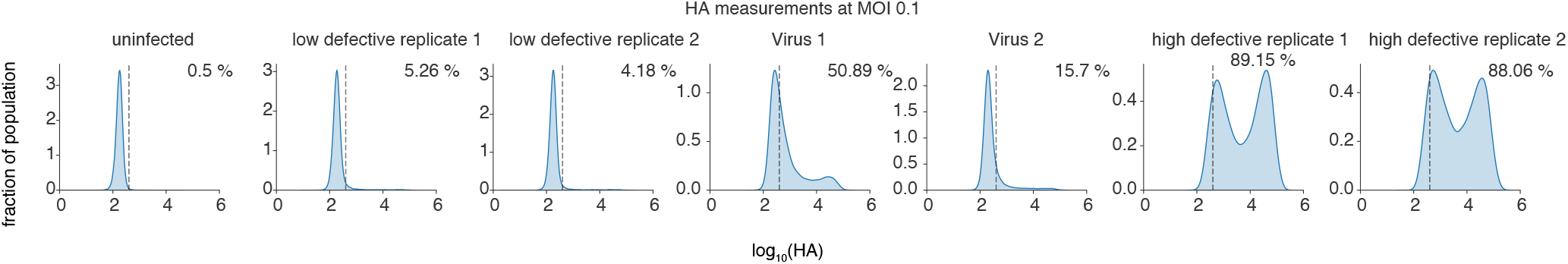
Analysis of influenza populations used in **Figure 4**A,B, and C. A549 cells were infected with the indicated populations, and, at 9h post-infection, stained with a monoclonal antibody against viral HA protein. Virus 1 and Virus 2 were the two populations used in experiments in **Figure 4**. Populations were grown as described in Materials and Methods. The dotted line indicates the gate at which positivity was called, in this case the 99.5^th^ percentile of the uninfected control.

**Figure 4–Figure supplement 2.**
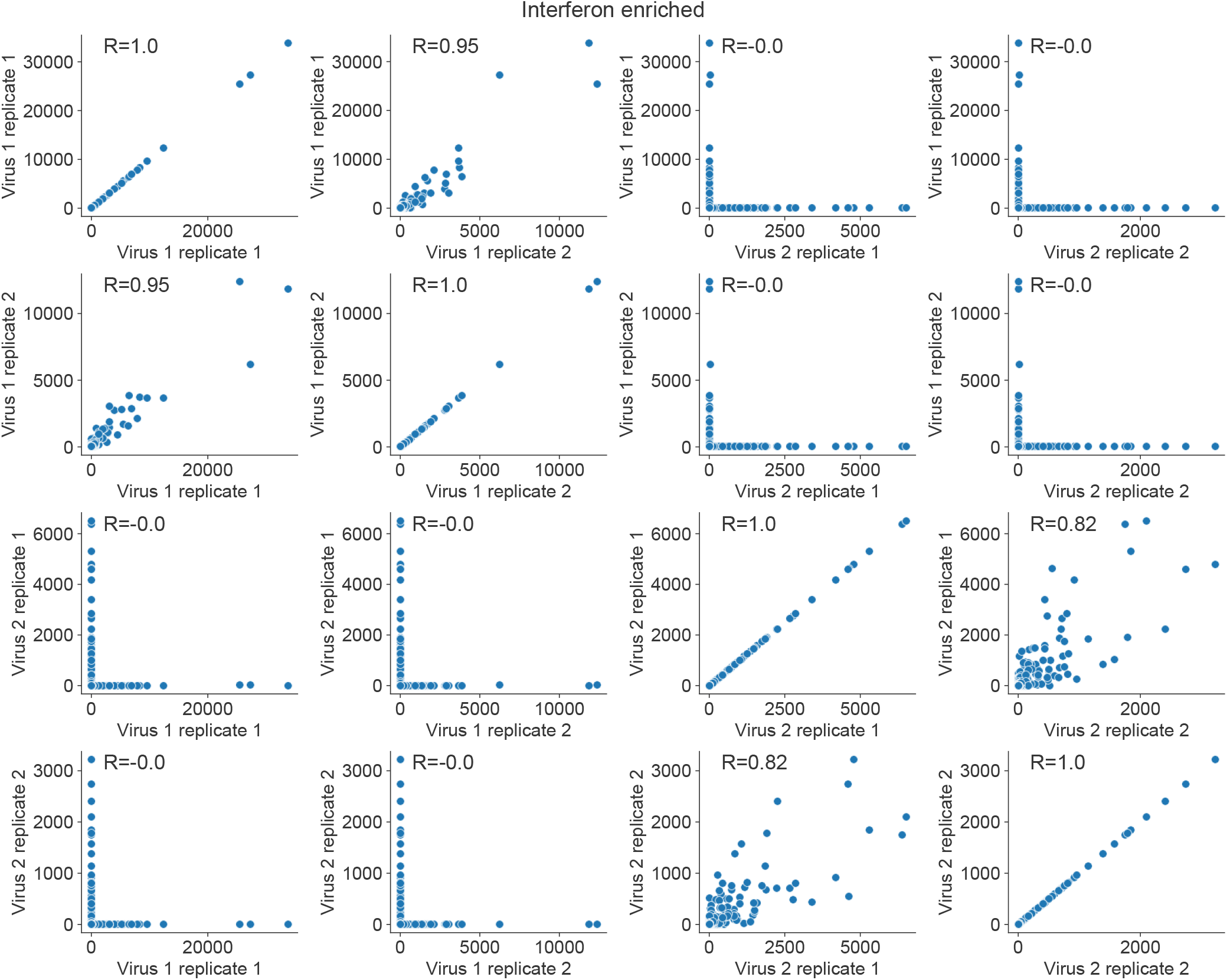
Sampling stochasticity for interferon-enriched samples for **Figure 4**B. Numbers represent raw counts of deletion-spanning junctions. Each point is a unique deletion. Virus 1 and Virus 2 were the two biologically-independent populations analyzed for (**Figure 4**B).

**Figure 4–Figure supplement 3.**
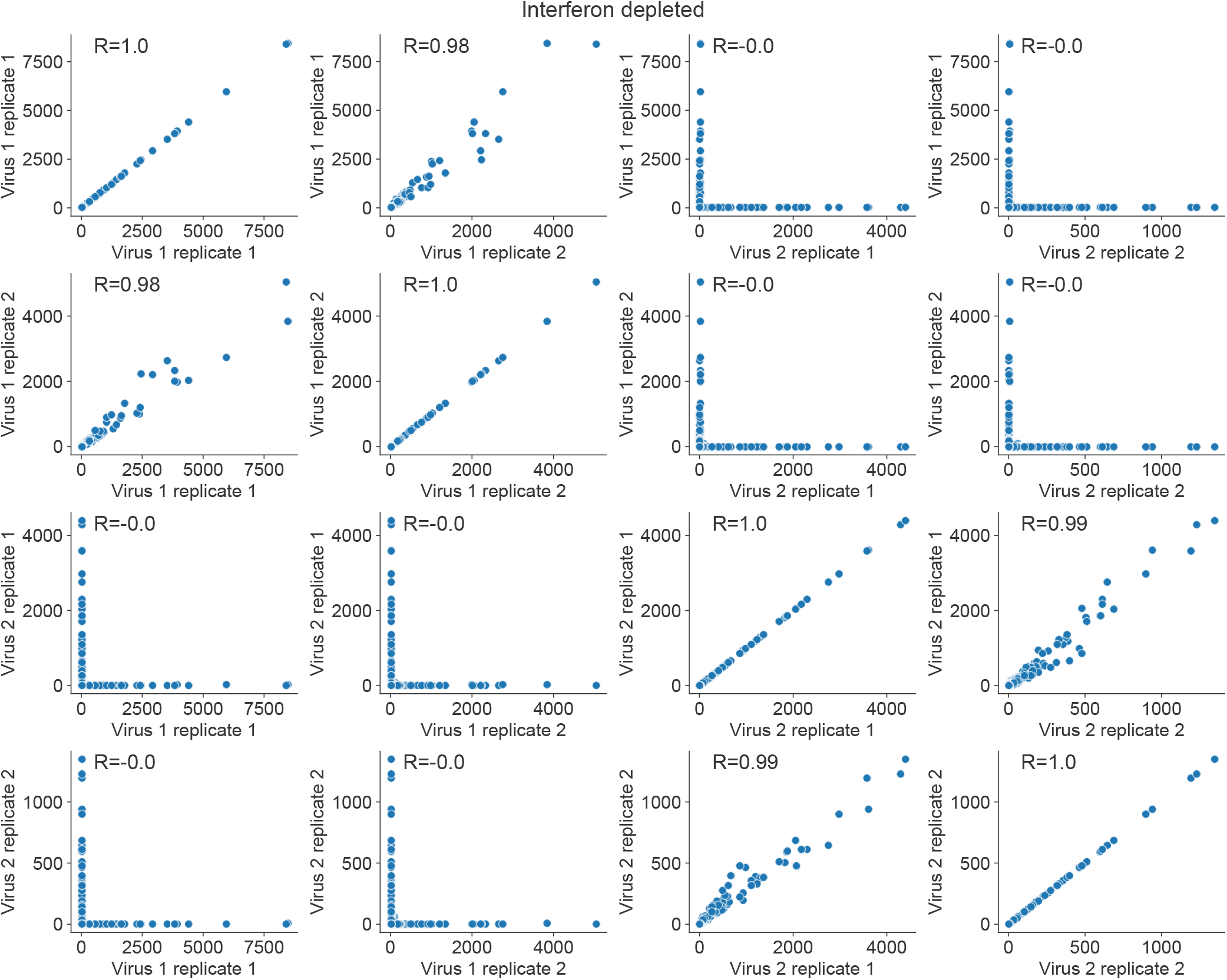
Sampling stochasticity for interferon-depleted samples for (**Figure 4**B). Numbers represent raw counts of deletion-spanning junctions. Each point is a unique deletion. Virus 1 and Virus 2 were the two biologically-independent populations analyzed for (**Figure 4**B).

**Figure 4–Figure supplement 4.**
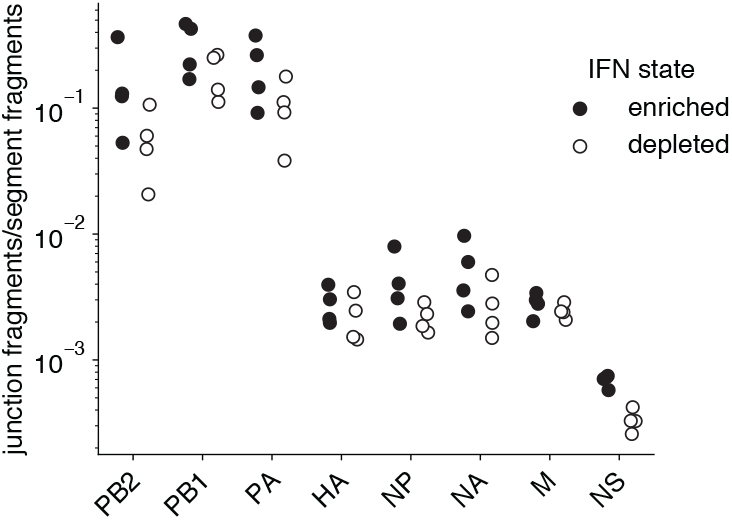
Raw junction data for **Figure 4**B. Total number of junction-spanning fragments normalized to the total number of fragments mapped per influenza genomic segment. n=4.

**Figure 4–Figure supplement 5.**
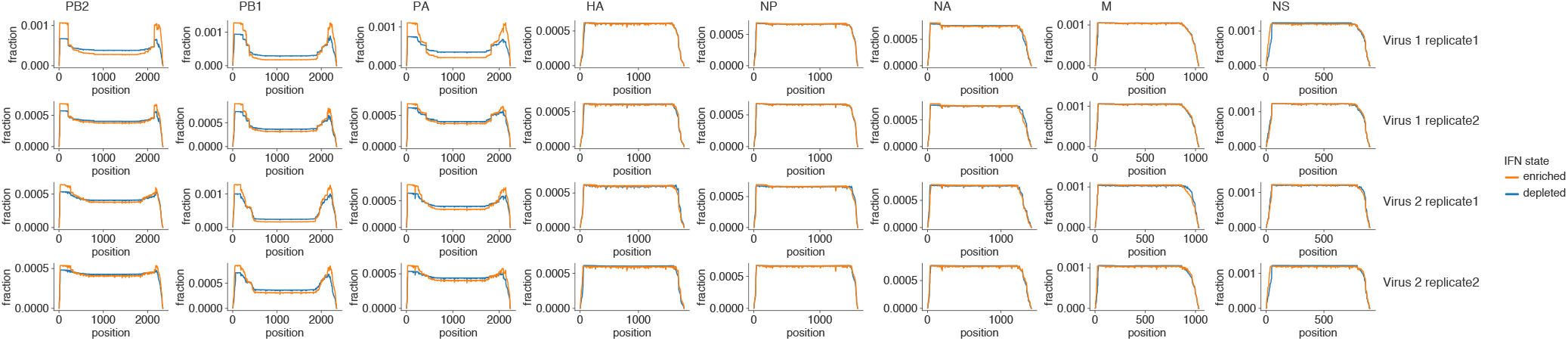
Raw mapping depth for sequencing data from **Figure 4**B. Depth per base corrected to fractional depth, area under each line sums to 1. Increased depth at 5’ and 3’ ends of polymerase segments consistent with increased fractions of deletions.

**Figure 4–Figure supplement 6.**
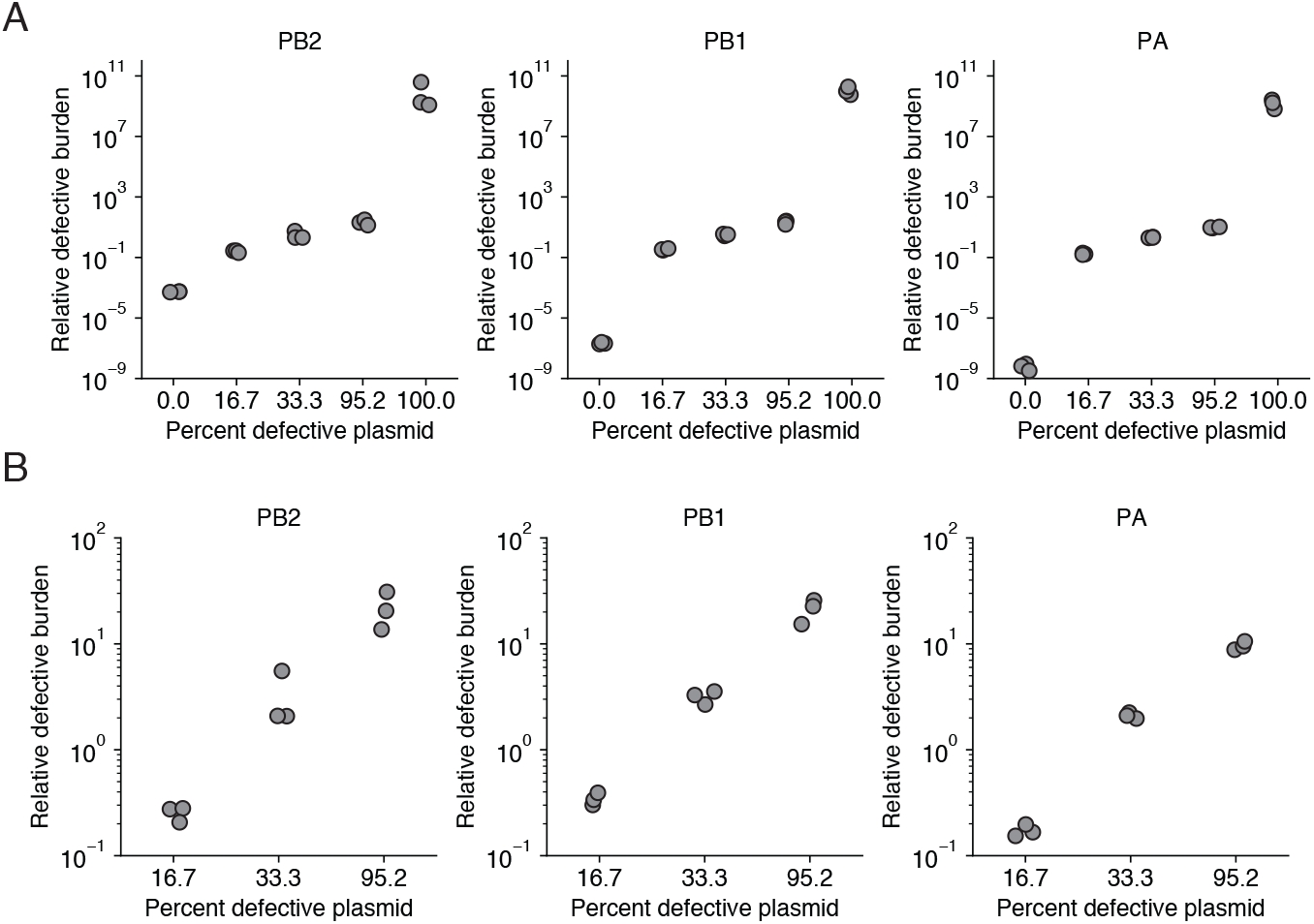
Validation of qPCR in **Figure 4**C. **(A)**Plasmids containing full-length PB2, PB1, and PA, and those containing PB2_316:267_, PB1_177:385_, and PA_328:137_ were mixed at the indicated molar ratios and 0.02 ng of the resultant mixture was analyzed by qPCR. Values represent the ratio of signal of our defective-spanning qPCR corrected for full-length-only signal. **(B)** Same as **(A)** but only intermediate values to show capacity to discriminate between extremes.

**Figure 4–Figure supplement 7.**
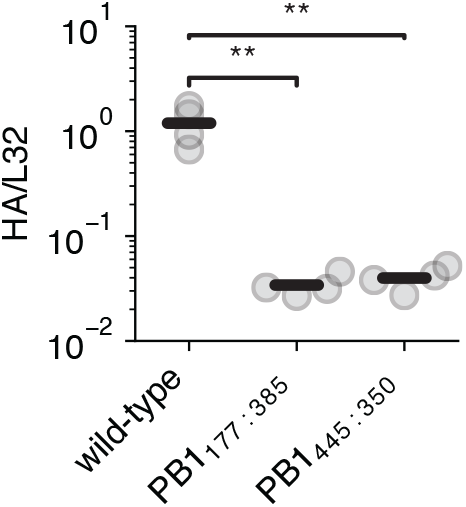
Validation of infections in **Figure 4**D, confirming levels of influenza transcript expression. qPCR measuring levels of HA transcript as normalized to the housekeeping control *L32* for the indicated influenza mutants infecting A549 cells at an MOI of 0.5 at 8h post-infection. The significant difference between PB1 deletion variants is not due to incorrect dosage, as shown by similar levels of influenza transcripts. Neither PB1 mutant is replication-competent, as shown by the reduced influenza transcript levels compared to a wild-type control. Asterisks represent significantly different values in all pairwise comparisons, two-tailed t-test, using Benjamini-Hochberg multiple-hypothesis correction at an FDR of 0.05. n=4.

**Figure 5–Figure supplement 1.**
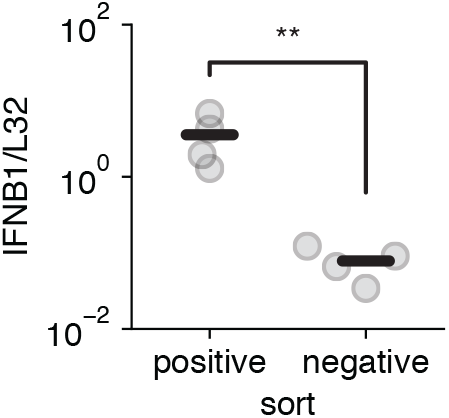
Validation of sort in **Figure 5**A. qPCR of sort in **Figure 4**A measuring levels of *IFNB1* transcript compared to the housekeeping control *L32*, for comparison with **Figure 5**A. Asterisks indicate significantly increased *IFNB1* transcript, two-tailed t-test p < 0.05. n=3.

**Figure 6–Figure supplement 1.**
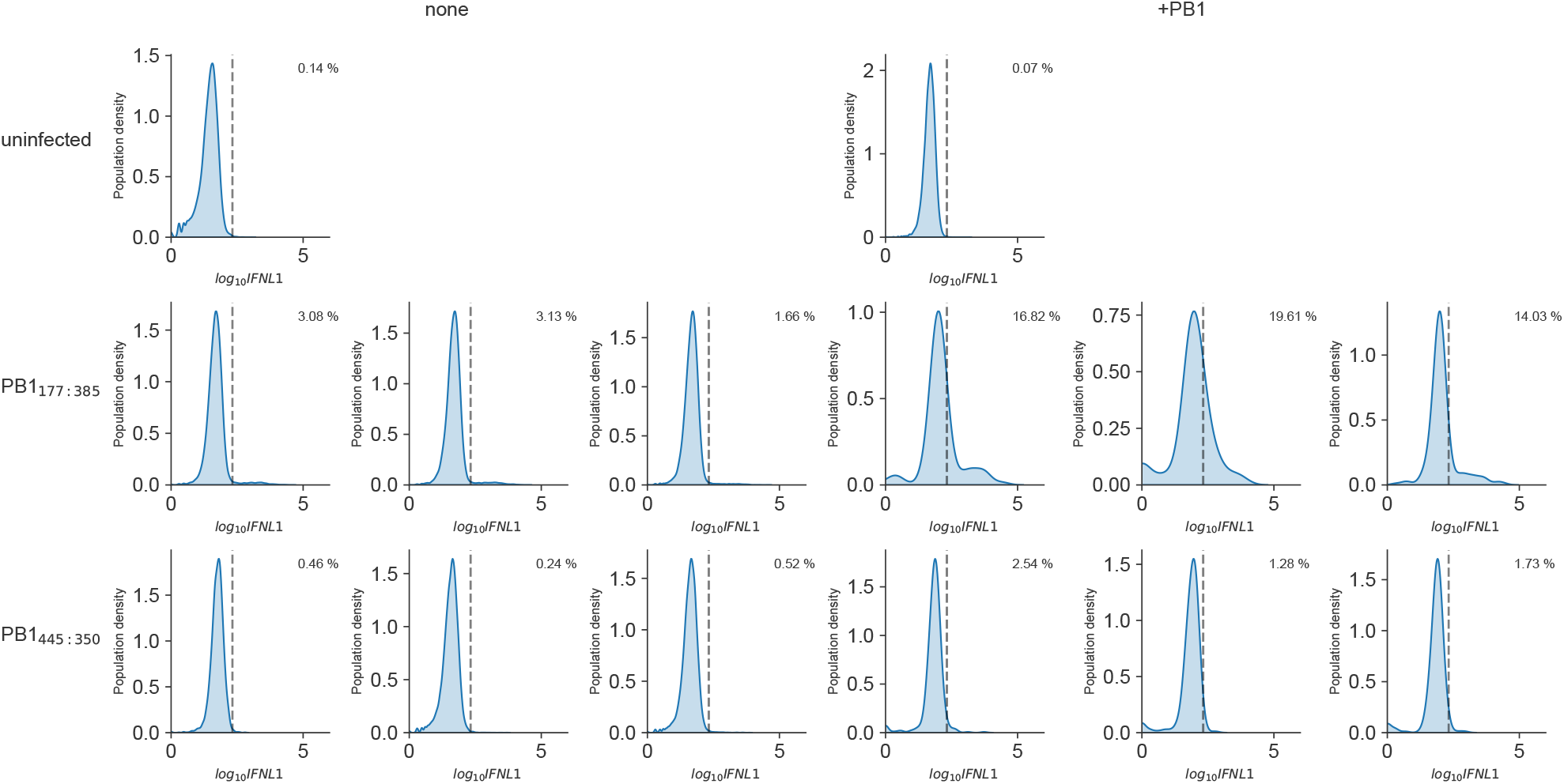
Gating for **Figure 6**B. Individual replicates shown. Dotted line represents positivity threshold Gating drawn on the uninfected control before application to infected samples, set at the 99.9^th^ percentile of uninfected. Data shown as kernel density estimates with Gaussian distributions. n=3.

**Figure 6–Figure supplement 2.**
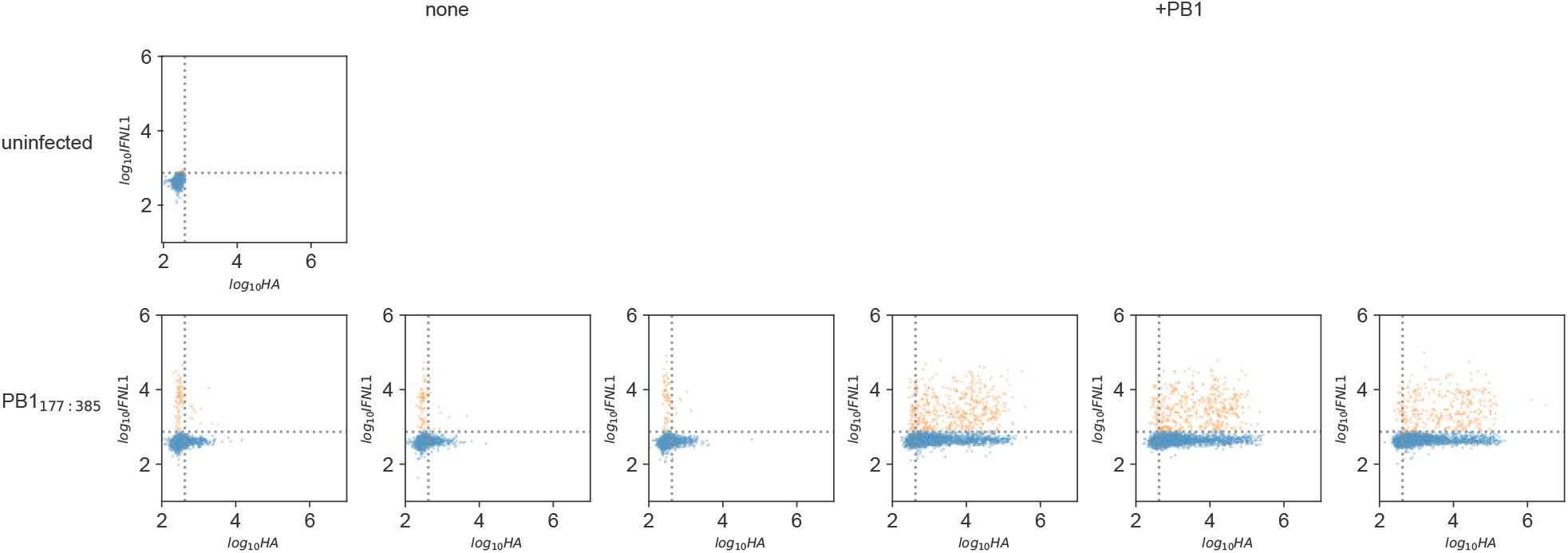
Gating for **Figure 6**C. Individual replicates shown. Dotted line represents gating. Gating drawn on the uninfected control before application to infected samples, set at the 99.9^th^ percentile of uninfected. Data were subsampled to 5000 events and shown as individual points. Points in orange were called as interferon-positive. n=3.

**Figure 7–Figure supplement 1.**
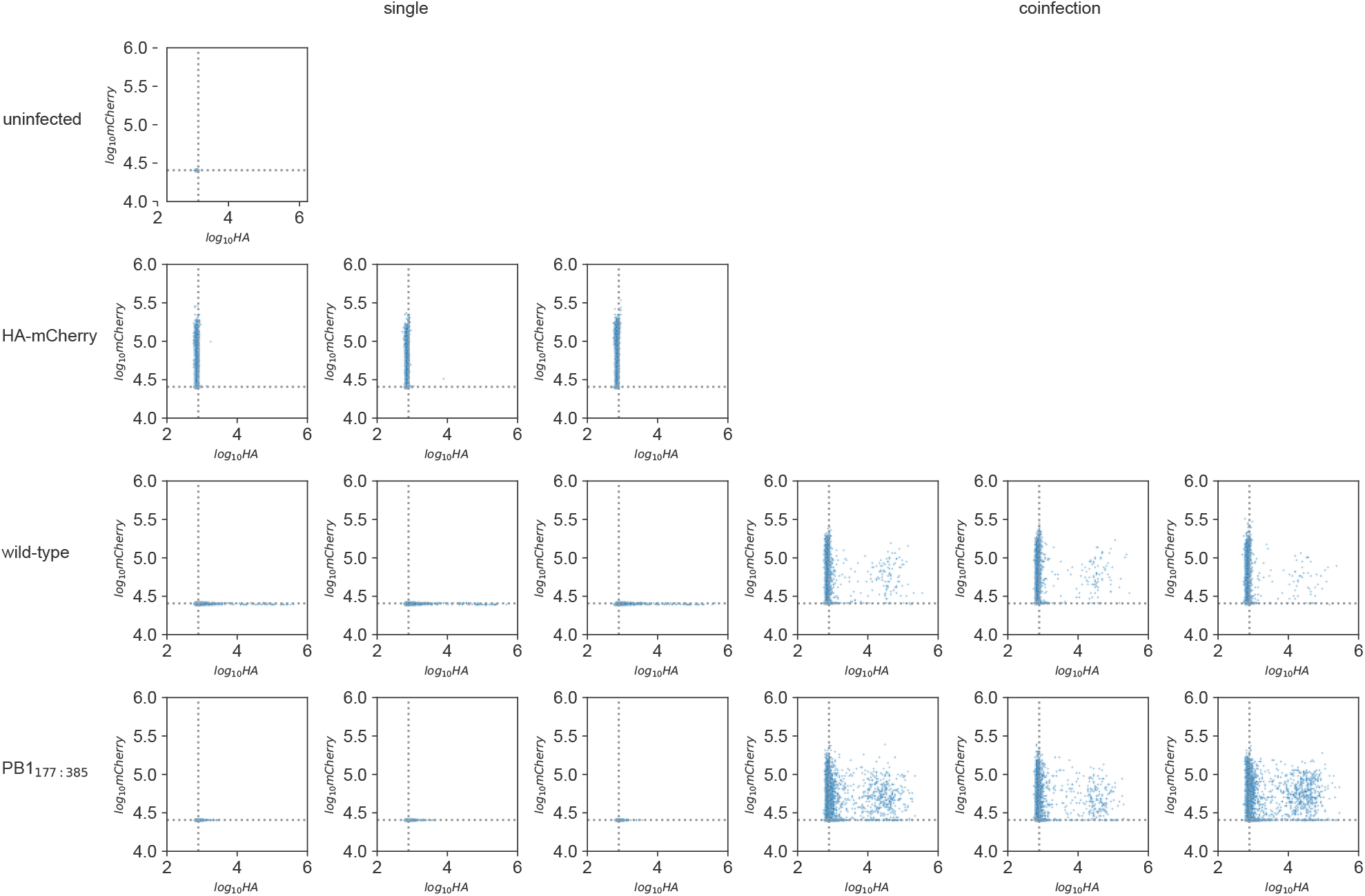
Gating for **Figure 7**A. Individual replicates shown. Dotted line represents gating. Gating drawn on the uninfected control before application to infected samples, set at the 99.95^th^ percentile of uninfected. Data were subsampled to 5000 events and shown as individual points. n=3.

**Figure 7–Figure supplement 2.**
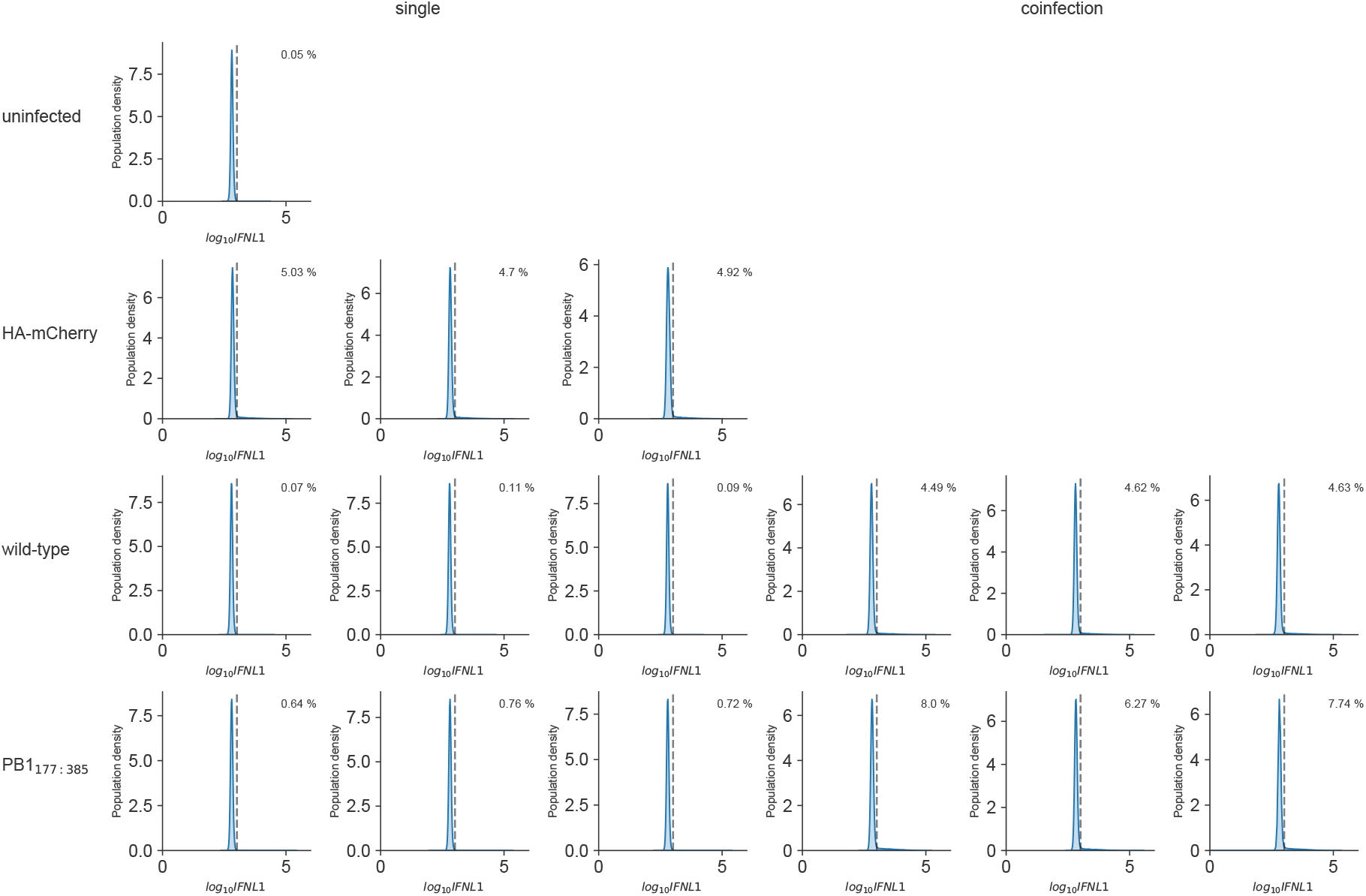
Gating for **Figure 7**A. Individual replicates shown. Dotted line represents gating. Gating drawn on the uninfected control before application to infected samples, set at the 99.95^th^ percentile of uninfected. Data shown as kernel density estimates with Gaussian distributions. n=3.

**Figure 7–Figure supplement 3.**
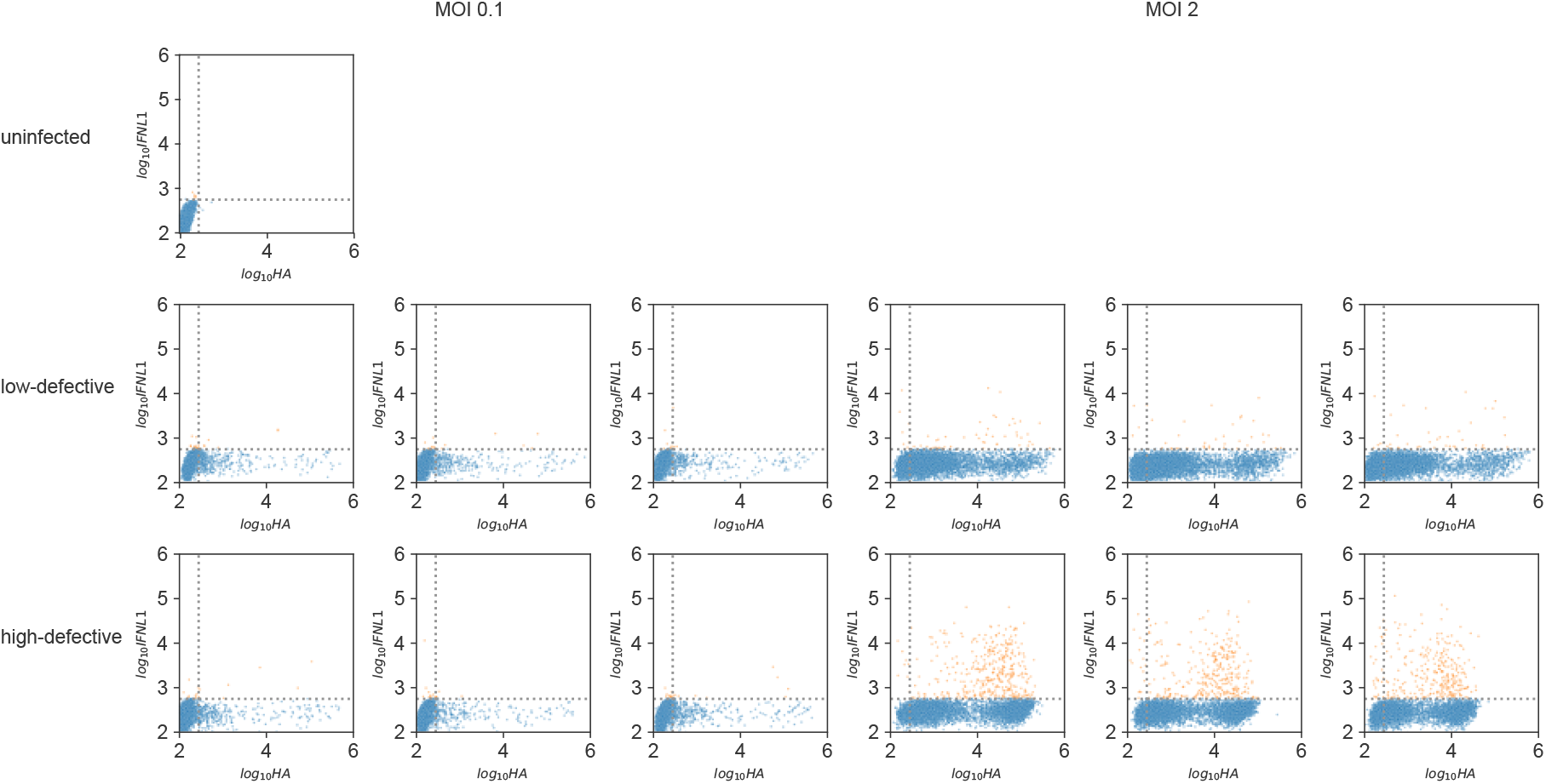
Gating for**Figure 7**B. Individual replicates shown. Dotted line represents gating. Gating drawn on the uninfected control before application to infected samples, set at the 99.9^th^ percentile of uninfected. Data were subsampled to 5000 events and shown as individual points. Points in orange were called as interferon-positive. n=3.

